# The development of brain pericytes requires expression of the transcription factor *nkx3.1* in intermediate precursors

**DOI:** 10.1101/2023.06.21.545941

**Authors:** Suchit Ahuja, Cynthia Adjekukor, Qing Li, Katrinka M. Kocha, Nicole Rosin, Elodie Labit, Sarthak Sinha, Ankita Narang, Quan Long, Jeff Biernaskie, Peng Huang, Sarah J. Childs

## Abstract

Brain pericytes are one of the critical cell types that regulate endothelial barrier function and activity, thus ensuring adequate blood flow to the brain. The genetic pathways guiding undifferentiated cells into mature pericytes are not well understood. We show here that pericyte precursor populations from both neural crest and head mesoderm of zebrafish express the transcription factor *nkx3.1* develop into brain pericytes. We identify the gene signature of these precursors, and show that an *nkx3.1, foxf2a*, and *cxcl12b* -expressing pericyte precursor population is present around the basilar artery prior to artery formation and pericyte recruitment. The precursors later spread throughout the brain and differentiate to express canonical pericyte markers. Cxcl12b-Cxcr4 signaling is required for pericyte attachment and differentiation. Further, both *nkx3.1* and *cxcl12*b are necessary and sufficient in regulating pericyte number as loss inhibits and gain increases pericyte number. Through genetic experiments we have defined a precursor population for brain pericytes and identified genes critical for their differentiation.

## Introduction

Endothelial cells and pericytes are key partners in the brain microvessel network. Endothelial cells line the luminal side of vessels, and pericytes attach to the abluminal side of endothelial cells. Pericytes stabilize microvessels by laying extracellular matrix around endothelial cells and regulating vascular tone (1–3). In addition, pericytes regulate postnatal endothelial sprouting and endothelial morphogenesis via VEGF and TGF-β signaling respectively (2, 4). Consequently, pathologies involving cranial hemorrhage, vessel dilation, and vessel structural defects are common when pericytes are reduced or absent due to loss of key pericyte signalling pathways, Pdgfb (5–9). Notch activity is also critical for emergence of perivascular mural cells, particularly smooth muscle cells (10–12). Loss- and gain-of-function of Notch3 receptor in zebrafish revealed a role in brain pericyte proliferation (13). In mice, pericyte loss due to Notch signaling deficiency leads to arteriovenous malformations (14).

Owing to its critical function in vascular homeostasis, pericyte development has received much attention. Quail-chick chimeras showed that pericytes of the forebrain are neural crest derived while aortic pericytes originate from Pax1+ and FoxC2+ sclerotome (15, 16). Mesodermal and neural crest origins of pericytes have also been shown in the zebrafish, where pericytes of the anterior midbrain originate from neural crest and those in the hindbrain and trunk are derived from the paraxial mesoderm (17). Transcriptional and signalling pathways that promote pericyte differentiation include the forkhead box transcription factors FoxC1 and FoxF2 that are required for brain pericyte differentiation and blood-brain barrier maintenance (18, 19). Furthermore, mice and zebrafish lacking FoxC1 and FoxF2 show cerebral hemorrhages (18–20). In line with this, humans with risk loci near *FOXF2* are more susceptible to stroke and cerebral small vessel disease (21).

While pathways promoting pericyte differentiation have been discovered, the initial signals triggering the convergent differentiation of brain pericyte precursors from two different germ layers are unknown. Here we describe the role of a homeobox transcription factor Nkx3.1, which is required in pericyte precursors originating in both neural crest and paraxial mesoderm. *nkx3.1^−/−^* mutants exhibit fewer pericytes on brain vessels and brain hemorrhage. Single cell sequencing on *nkx3.1*-lineage cells reveals that pericyte precursors are marked by the transcription factors *tbx18* and *foxf2a* amongst other genes. Furthermore, we show that chemokine ligand *cxcl12b/sdf1* is expressed in pericyte precursors and functions downstream of *nkx3.1* during pericyte development. Taken together, our study defines a previously unknown pericyte precursor population and a novel Nkx3.1-Cxcl12b cascade during pericyte development.

## Results

### Brain pericytes originate from a lineage marked by *nkx3.1*

An early marker of the zebrafish sclerotome is the transcription factor, *nkx3.1*. Trunk pericytes are derived from *nkx3.1*-expressing sclerotome precursors, although *nkx3.1* is downregulated when trunk pericytes differentiate (16, 22). Precursor markers for brain pericytes have not yet been identified. Brain pericytes form from two germ layers (neural crest and mesoderm) and an understanding of the convergent genetic program to differentiate cells from different origins into pericytes is also lacking. *nkx3.1* is expressed in the ventral head mesenchyme and trunk of the developing embryo at 16 and 30 hpf, as shown by in situ hybridization (Fig. 1A-B, arrowheads). *nkx3.1* expression in ventral head mesenchyme is still present at 30 hpf but greatly reduced. It is undetectable by 48 hpf (Fig. 1C). Expression of *nkx3.1* occurs far earlier than that of the pericyte marker pdgfrβ, first expressed at 48 hpf in the basilar artery(13). To determine whether brain pericytes originate from *nkx3.1* lineage, we made use of the transgenic lines *TgBAC(nkx3.1:Gal4)^ca101^* and *Tg(UAS:NTR-mCherry)^c264^* to raise *nkx3.1:Gal4; UAS:Nitroreductase-mCherry*, hereafter known as *nkx3.1^NTR-mcherry^*embryos where *nkx3.1* lineage cells are labelled with mCherry. mCherry perdures after native *nkx3.1* mRNA is downregulated, allowing us to track cell lineage beyond the time that *nkx3.1* mRNA is normally expressed (22–24). At 4 days post fertilization (dpf), we observe *nkx3.1^NTR-mcherry^* cells in the perivascular zone surrounding endothelial cells (Fig. 1D-E’, arrowheads). These *nkx3.1*-lineage cells show a pericyte-like morphology with a round soma and processes that wrap around endothelial cells (Fig. 1E-F’, arrowheads). Furthermore, we found a complete overlap in expression between *nkx3.1^NTR-mcherry^* and a transgenic reporter of pericytes, *TgBAC(pdgfrb:GFP)^ca41^* at 75 hpf (Fig. 1G-I). This confirms that *nkx3.1^NTR-mcherry^* perivascular cells in the brain are pericytes at 75 hpf. Since *nkx3.1* is expressed prior to *pdgfr*β, but their later expression is completely overlapping, this data suggests that *nkx3.1* is expressed in pericyte precursors. To characterize the cellular behavior of *nkx3.1^NTR-mcherry^* perivascular cells, we imaged labelled cells in the embryonic brain from 55-65 hpf. Timelapse reveals that *nkx3.1^NTR-mcherry^* perivascular cells migrate and proliferate on blood vessels (Fig.1J-M, white and yellow arrowheads; S1 Movie), similar to previously described cellular behavior of pericytes (17).

**Fig. 1:**
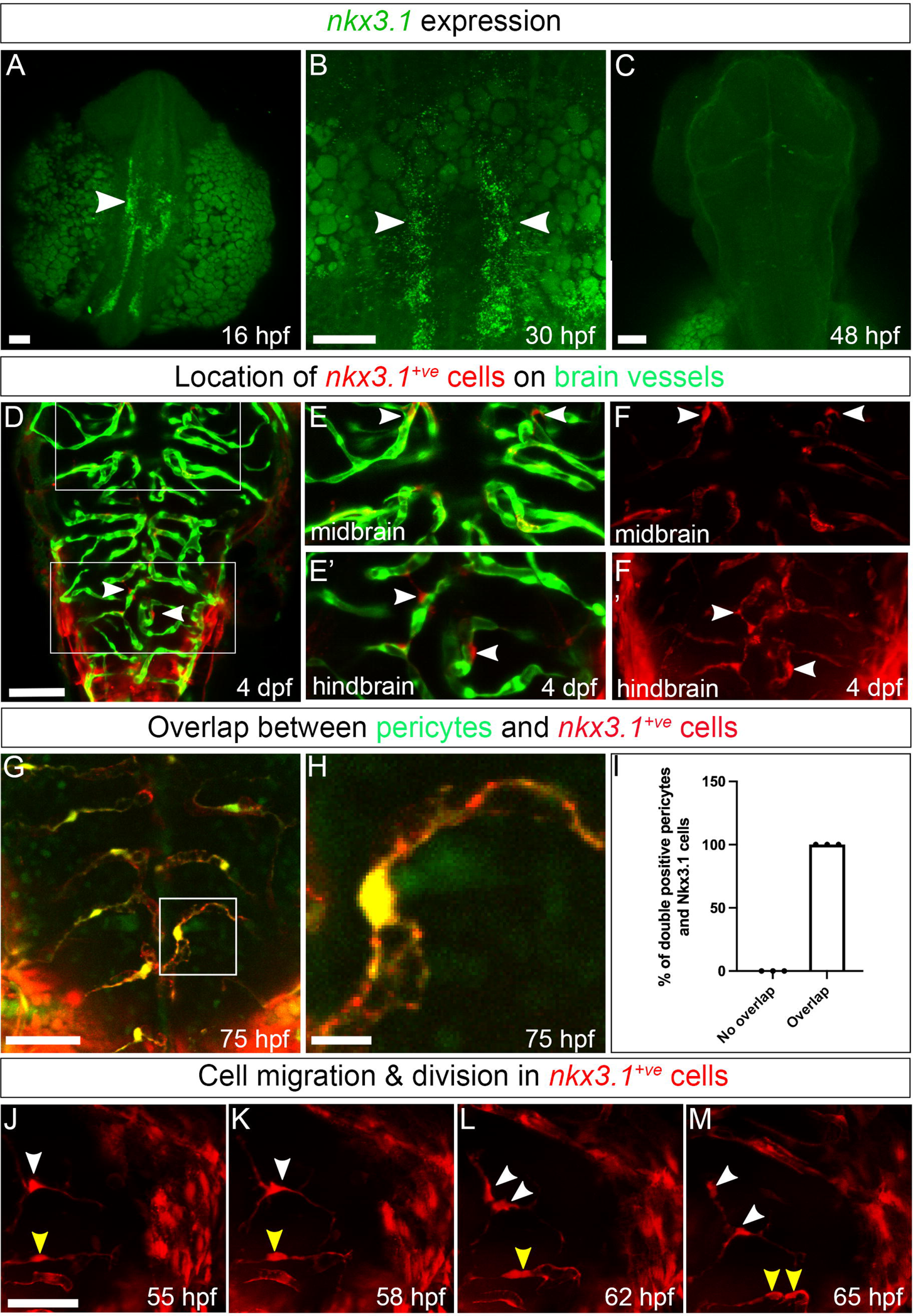
*nkx3.1* expression patterns and cell behavior. All images are captured dorsally and the anterior (A) and posterior (P) axis is marked. (A-C) Expression of *nkx3.1* by HCR in situ hybridization. (A) At 16 hpf, *nkx3.1* is expressed in the hindbrain and anterior trunk. Arrowheads mark *nkx3.1* expression. (B) At 30 hpf, *nkx3.1* is expressed in the posterior head and trunk. (C) At 48 hpf, *nkx3.1* expression is not detectable. (D-E’) *nkx3.1^NTR-mcherry^* cells (red) are adjacent to endothelium (green; *Tg(flk:GFP)*) in brain vessels at 4 dpf. Pericytes in midbrain (arrowheads, E, F) and hindbrain (arrowheads, E,’ F’) are denoted in dual channel (E, E’) and single channel pericyte (F, F’) images. (G) Brain pericytes labelled by *TgBAC(pdgfr*β*:GFP)* co-express *nkx3.1^NTR-mcherry^* 75 hpf. (H) Enlargement of an individual brain pericyte marked by a square in G. (I) Quantification brain pericytes co-expressing Nkx3.1 at 75 hpf (N=3 experiments and 30 embryos). (J-M) Single images from timelapse of *nkx3.1^NTR-mcherry^* cells in the midbrain. White and yellow arrowheads track individual cells that migrate and divide with time. Scale bar in all images is 50µm.

Previous lineage-tracing data suggests that pericytes in the zebrafish hindbrain originate from mesoderm and that midbrain pericytes originate from neural crest (17). Pericytes in both hindbrain and midbrain express the *nkx3.1* transgene (Fig. 1D). We next used lineage tracing of mesoderm and/or neural crest progenitors to test whether *nkx3.1*-expressing cells arose from one or both germ layers. We used Cre drivers for mesoderm *Tg(tbx6:cre;myl7:GFP)* or neural crest *Tg(sox10:cre;myl7:GFP)* together with a floxed reporter *Tg(loxp-stop-loxp-H2B-GFP*) to lineage label mesodermal or neural crest progeny, respectively. We crossed these fish with an *nkx3.1* reporter *TgBAC(nkx3.1:Gal4)*, pericyte reporter *TgBAC(pdgfrb:Gal4FF)*, or endothelial reporter *Tg(kdrl:mCherry)*. This strategy labels cells expressing *pdgfr*β, *nkx3.1* or *kdrl* in red and mesodermal or neural crest derivatives as green (depending on the experiment). We imaged double positive cells to identify their lineage and observe that both mesoderm and neural crest progenitors contribute to both hindbrain and midbrain pericytes by lineage tracing either pdgfrβ or *nkx3.1* (S1 Fig.).

Taken together, our data shows that *pdgfr*β*-*expressing brain pericytes originate from a *nkx3.1* positive precursor, which in turn is generated from *tbx6 and sox10 expressing* mesodermal and neural crest progenitors. *nkx3.1* is therefore a transcription factor expressed in precursors of brain pericytes, but not in mature brain pericytes.

### Nkx3.1 function is required for brain pericyte development

To determine whether Nkx3.1 function is necessary for brain pericyte development, we made *nkx3.1* mutant zebrafish using CRISPR-Cas9. *nkx3.1^ca116^* mutants have a 13 bp deletion in exon 2 that is predicted to lead to premature stop prior to the homeobox domain (S2 Fig.). *nkx3.1* ^−/−^ mutant embryos show no gross morphological defects and adults are homozygous viable, although with reduced lifespan of approximately 1 year. At 75 hpf, we observed no difference in total pericyte number (sum of mid- and hindbrain pericytes) between *nkx3.1^−/−^* mutants and their heterozygous and wildtype siblings (S3 Fig.), however a previous study showed that *nkx3.1* transcripts are maternally contributed to the developing embryo (25). To remove this maternal contribution, we crossed *nkx3.1^+/-^* males with *nkx3.1^-/-^* females. Maternal zygotic (MZ) *nkx3.1^−/−^*embryos show a strong phenotype including brain hemorrhage and hydrocephalus at 52 hpf as compared to controls (Fig. 2A-B). The average percentage of MZ *nkx3.1^−/−^* embryos exhibiting brain hemorrhage at 52 hpf (54%) was significantly higher as compared to *nkx3.1^+/-^* siblings (12.5%; Fig. 2C). Not surprisingly, given altered hemodynamics after hemorrhage, central artery (CtA) vessel diameter was decreased from 5.9 to 5.4 µm on average (Fig 2D).

**Fig. 2:**
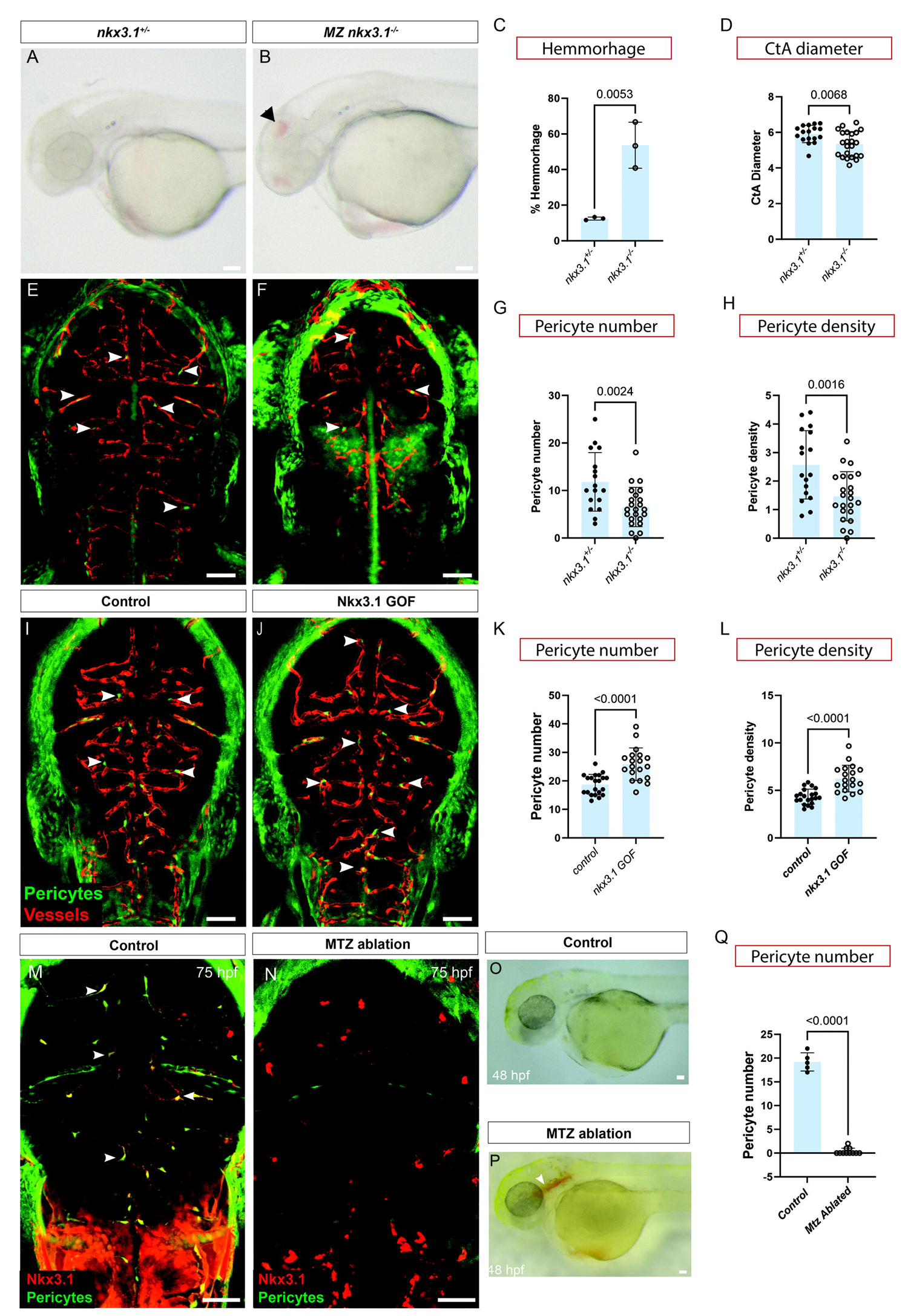
Nkx3.1 function is required to regulate brain pericyte numbers. Lateral view of control *nkx3.1*^+/-^ (A) and *nkx3.1*^−/−^ MZ (B) mutants showing brain hemorrhage (arrowhead) at 52 hpf, and quantification (C; N=3, proportions of hemorrhage). (D) *nkx3.1*^−/−^ have decreased CtA vessel diameter in comparison to *nkx3.1*^+/-^ hets. Dorsal images of *nkx3.1*^+/-^ hets (E) and *nkx3.1*^−/−^ (F) embryos expressing *Tg(pdgfr*β*:GFP)* and *Tg(kdrl:mCherr*y) showing fewer brain pericytes (arrows) in mutants at 75 hpf. Quantification of (G) decreased pericyte number and (H), decreased pericyte density (defined as the number of pericytes divided by the length of the vessel network) in mutants. In comparison to control wildtype embryos (I*, Tg(hsp70l:tBFP,*), *nkx3.1* gain-of-function (GOF) embryos expressing *Tg(hsp70l:tBFP-2a-nkx3.1))* show more pericytes (J, arrowheads) ( n=20 control and 19 GOF)) as quantified (K). Pericyte density is also increased (L). Dorsal views of embryos labelled with *TgBAC(nkx3.1:Gal4)* and *Tg(UAS:NTR-mCherry)* under fluorescence (M-N) or under brightfield (O-P) that are untreated (M,O) or treated with metronidazole (N,P) to ablate *nkx3.1*-expressing cells. Ablated embryos show brain hemorrhage (P, arrowhead) at 48 hpf (P). (Q) Quantification of pericyte number in ablated embryos. Statistical significance was calculated using the Student’s t-test (n=5 wildtype and 11 *nkx3.1* mutants). Scale bars are 50µm. The data underlying this figure can be found in S3 Table.

Importantly, MZ *nkx3.1^−/−^* showed significantly fewer pericytes at 75 hpf, as observed by confocal microscopy (Fig. 2E-G). This is consistent with the brain hemorrhage phenotype, which is a reported consequence of lack of pericytes (26). Furthermore, the density of pericytes (defined as the number of pericytes divided by the length of the vessel network) is also reduced (Fig 2H). The total length of the Central Arteries (CtA network length) is unchanged suggesting that the endothelial patterning is unaffected (S4 Fig.) These phenotypes are maintained at into early larval stages; both pericyte number and pericyte density are also decreased at 5dpf in *nkx3.1* mutants, while CtA network length is unchanged (S4-S5 Figs.). All work further therefore used maternal-zygotic (MZ) *nkx3.1* mutants.

To test the sufficiency of Nkx3.1 for pericyte development, we made an *nkx3.1* gain-of-function (GOF) transgenic line *Tg(hsp70l:tBFP-2a-nkx3.1)* expressing Blue fluorescent protein (BFP) fused with *nkx3.1*. *Tg(hsp70l:tBFP)* as a control. Overexpression of *nkx3.1* using heat shock from 29-30 hpf, when first brain pericytes are emerging, results in significantly more brain pericytes at 75 hpf, compared to expression of tBFP alone (Fig. 2I-K). Brain vessels of *nkx3.1* GOF embryos appear grossly morphologically normal. Furthermore, the density of pericytes on vessels is increased in *nkx3.1 GOF* suggesting the increased number of pericytes is spread more tightly on the same length of vessel (Fig 2L). Thus *nkx3.1* is both necessary and sufficient for brain pericyte development.

As a second method to test the necessity of *nkx3.1* precursors, we made used the *nkx3.1^NTR-mcherry^*transgenic line to ablate *nkx3.1^NTR-mcherry^* cells by treating with 5mM metronidazole from 24-48 hpf, a time window during which brain pericyte differentiation is occurring (17). Consistent with the MZ *nkx3.1^−/−^* phenotype, ablation of *nkx3.1^NTR-mcherry^*positive cells in transgenic embryos treated with metronidazole showed no pericytes at 75 hpf. *nkx3.1* thus appears to be expressed in all brain pericyte precursors as no pericytes remained after ablation (Fig2 M-N, Q). We also observed brain hemorrhage at 48 hpf after ablation (Fig2 O-P).

### Pericyte precursors share markers with fibroblasts

Although pericytes derive from *tbx6*-positive mesodermal cells and *sox10*-positive neural crest cells, intermediate progenitors have not yet been defined. We have shown that *nkx3.1*-expressing cells differentiate into pericytes and that *nkx3.1* is a marker of pericyte progenitors. Therefore, *nkx3.1* positive cells sampled at a stage prior to pericyte differentiation are a unique population to interrogate the gene expression profile of pericyte precursors. Since not all *nkx3.1* positive cells become brain pericytes (i.e., some become fibroblasts derived from sclerotome cells of the trunk and other lineages), embryos expressing the *nkx3.1^NTR-mcherry^*transgene were dissociated from wildtype 30 hpf zebrafish embryos and mCherry-positive FACs sorted cells were subjected to single cell RNA sequencing (scRNAseq; Fig. 3A, B).

**Fig. 3:**
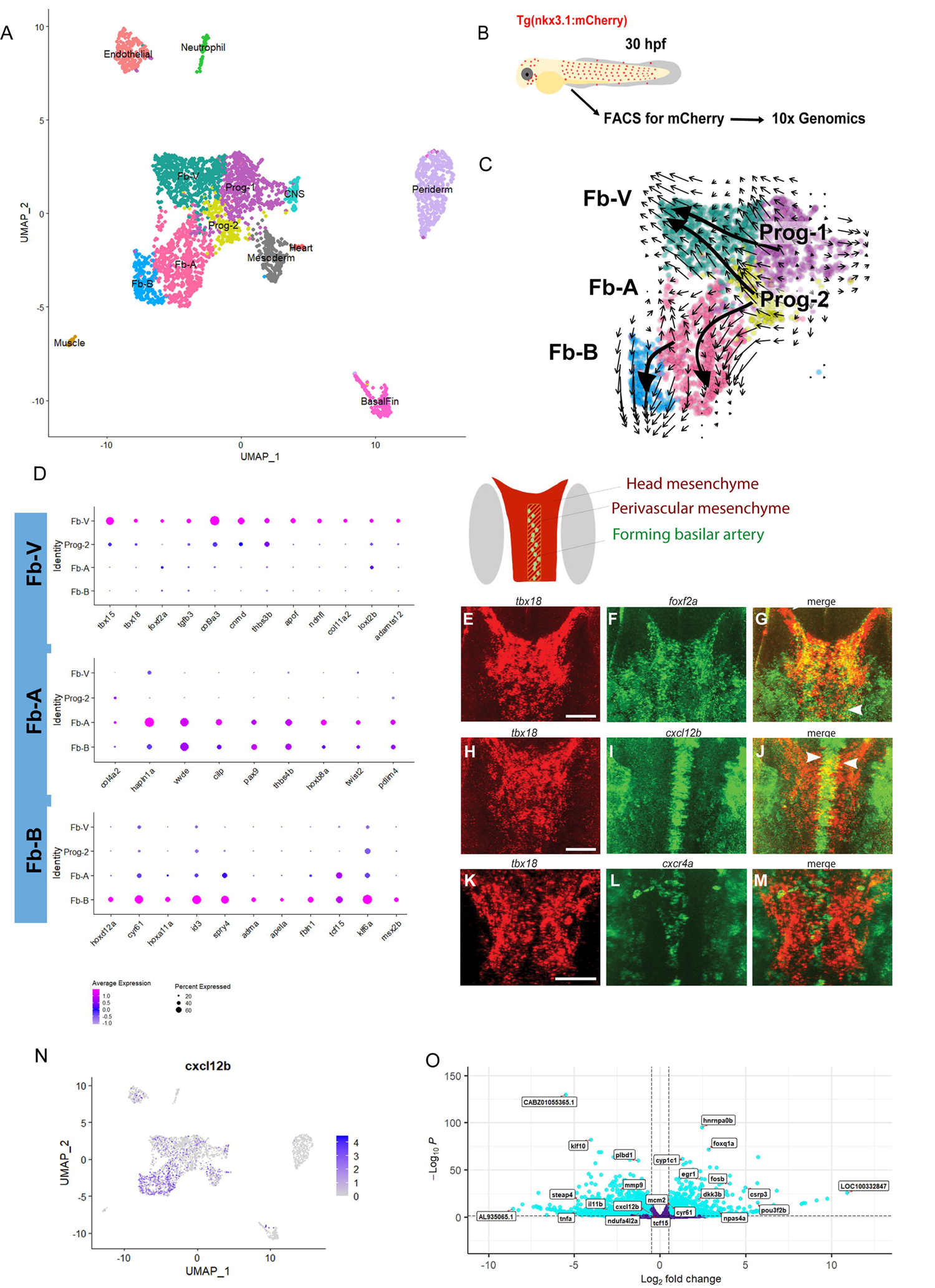
Next generation sequencing analysis of *nkx3.1^NTR-mcherry^* and *nkx3.1^−/−^* embryos at 30 hpf. (A) Single cell clusters of *nkx3.1^NTR-mcherry^* embryos at 30 hpf. (B) Schematic showing workflow for single cell sequencing of *nkx3.1^NTR-mcherry^*cells. (C) RNA-velocity analysis of subclustered *nkx3.1^NTR-mcherry^*cells of the Progenitor and FB-V, FB-A, and Fb-B clusters. (D) Dot plots showing key marker genes of FB-V, FB-A, and Fb-B clusters. (E-M) HCR expression analysis of Fb-V genes at 36 hpf with the schematic showing relative locations of the ventral head mesenchyme and the forming basilar artery in a dorsal schematic of the whole zebrafish brain (grey marks the position of the eyes). (E-M) HCR fluorescent in situ hybridization imaged by confocal showing sub-stacks in the region of the forming basilar artery and precursor area. (E-G) *tbx18* (red) and *foxf2a* (green) show expression overlap at 36 hpf in the ventral head (yellow, arrowheads). (H-J) *tbx18* (red) and *cxcl12b* (green) show expression overlap at 36 hpf in the ventral head (yellow, arrowheads). (K-M) *tbx18* (red) is expressed in the perivascular space surrounding the *cxcr4a* (green) expressing basilar artery in the ventral head at 36 hpf. (N) *cxcl12b* feature plot showing its expression across clusters including Fb-V. (O) Bulk sequencing volcano plot of *nkx3.1^−/−^* embryos at 30 hpf. Scale bar is 50µm.

3359 cells obtained from two biologically independent samples passed quality control analysis. Using the Uniform Manifold Approximation and Projection algorithm (UMAP) in Seurat (27), we detected thirteen different populations (Fig. 3A, S1 Table). We identified unique markers in each cluster (Fig. 3D, S8-11 Fig.). Cluster assignment used comparisons with published scRNAseq data (S1 Table) (28–32). Canonical fibroblast markers (*col1a1a, col1a1b, pdgfr*α*, col5a1, mmp2;* S7 Fig) are expressed by a group of connected clusters, including three more differentiated clusters (Fb-A, Fb-b, Fb-V) and one cluster that expresses both fibroblast and mitotic markers (Progenitor-like; Prog-2). Additional clusters not followed up here express mesoderm and heart, CNS, endothelial, Basal fin (fibroblasts of the epithelial layer), CNS, or neutrophil markers.

To understand the relationships between clusters, we used RNA velocity which measures transient transcriptional dynamics to infer the differentiation process, from analysis of RNA splicing information in sequencing data. We subclustered 2120 cells in 5 clusters (Prog-1/2 and Fb-A/B/V; Fig 3C). Using RNA velocity, we find that flow direction is consistent with two progenitor pools feeding into the fibroblast clusters, with the Progenitor 1 (Prog-1) cluster feeding into Fb-V and Progenitor 2 (Prog-2) feeding into all 3 fibroblast-like clusters (Fb-A, Fb-B and Fb-V). Fb-B is a smaller cluster (6% of sorted *nkx3.1* cells) that appears to arise from the Fb-A fibroblast.

Based on a gene expression profile overlapping with some pericyte markers, the Fb-V cluster most likely represents a pericyte progenitor cluster. Canonical pericyte markers like *pdgfr*β*, cspg4* and *notch3* are present in scattered cells in these clusters at this time point, but not yet enriched in Fb-V (S11 Fig.). However, an addition three pericyte markers, *tbx18*, *foxf2* and *cxcl12b* are enriched in Fb-V and mark *nkx3.1*-positive pericyte precursors, at this early stage prior to when canonical pericyte markers like *pdgfr*β are expressed.

*tbx18* is expressed in the embryonic zebrafish head paraxial mesoderm (33), and has enriched expression in adult mouse brain mural cells. A lineage trace of mouse *Tbx18* shows expression in both adult mouse pericytes and vascular smooth muscle cells (34, 35). *foxf2* is also enriched in the Fb-V cluster. We have previously shown that *foxf2a* and *foxf2b* are expressed in zebrafish brain pericytes and are essential for generating the proper number of brain pericytes (20, 36). Furthermore, mouse *Foxf2*, is enriched in, and critical for mouse brain pericyte formation (21, 31). A third gene we explore in the Fb-V cluster is *cxcl12b* (Fig. 3N). Vascular mural cells associated with the zebrafish coronary arteries and caudal fin vessels express *cxcl12b* (37, 38). In addition, mural cells of the mouse and human lung tissue express *Cxcl12* (39). Since our data is collected at a stage where there is scant information in other species about the differentiating pericyte gene expression profile and will be very useful for further studies (S1 Table).

To validate expression of genes of interest from the Fb-V cluster, we used in situ hybridization at 36 hpf when the first pericytes are associating with developing brain vessels. We find that *tbx18* is expressed in the perivascular space around the *cxcr4a+* basilar artery (Fig. 3K-M), where the first brain pericytes attach (17). We also detect *foxf2a* and *cxcl12b* expression in *tbx18+* cells (Fig. 3E-J). These data validate single cell sequencing results and confirm the presence of a perivascular cell type that co-expresses *tbx18*, *foxf2a*, and *cxcl12b* around the forming basilar artery, the first site of pericyte attachment (17).

The Fb-A and Fb-B clusters are diverging from Fb-V and express genes reminiscent of sclerotome including *pax9* and *twist2*, while the Fb-B cluster is enriched for fibroblast markers such as *cyr61,* and the *tcf15/paraxis*, involved in trunk mesoderm development. Both clusters express *thrombospondin 4b (thbs4b)*. Since Fb-B flows away from Fb-A, which flows away from Prog-2, this suggests that this lineage is differentiating into traditional fibroblasts that will go on to assume many different lineages, including those of the zebrafish trunk (23, 24).

### Loss of *nkx3.1* leads to transcriptional reduction of the chemokine *cxcl12*

To determine which genes within *nkx3.1*-expressing cells are important for their differentiation, we took advantage of *nkx3.1^ca116^* genetic mutant fish. At the identical stage to the scRNAseq (30 hpf), we sampled *nkx3.1* wildtype and MZ *nkx3.1* mutant embryos using bulk RNA sequencing. This revealed that 2788 genes were downregulated, and 2080 genes upregulated at 30 hpf (S2 Table). Among the genes downregulated at 30 hpf is *tcf15*, the pericyte marker *ndufa4l2a* and chemokine *cxcl12b* (Fig. 3O). Since we showed that *cxcl12b* is expressed in the pericyte progenitor cluster (Fb-V), we next interrogated the role of Cxcl12b in pericyte development.

### Cxcl12b signaling regulates brain pericyte number

Using in situ hybridization at 16 hpf and 24 hpf, we find that *cxcl12b* and *nkx3.1* are co-expressed in the ventral head mesenchyme of the developing embryo (S12 Fig.). We next tested expression of *cxcl12b* in *nkx3.1^−/−^* mutants. At 30 hpf, *cxcl12b* is strongly downregulated in the head of MZ *nkx3.1^−/−^* mutants at a location where the first pericytes will attach to the basilar artery (Fig. 4A-C). To test the role of Cxcr4-Cxcl12 signaling in pericyte development, we used AMD 3100, an inhibitor of the Cxcl12 receptor, Cxcr4. The number of brain pericytes and brain pericyte density is significantly reduced after treatment of zebrafish embryos with 100 µM AMD 3100 from 24-75 hpf (Fig. 4D-G), suggesting requirement of Cxcl12b signaling in pericyte development. The CtA network length is unchanged (S4 Fig). Consistent with this finding, gain of function induced by mRNA injection of 30 pg *cxcl12b* mRNA not only increases brain pericyte number in wildtype embryos but also increases brain pericyte number in MZ *nkx3.1* mutants at 75 hpf. However, GOF *cxcl12b* expression does not decrease hemorrhage when expressed in *nkx3.1* mutants (S13 Fig.). We note that there is no overlap in cxcr4a and cxcl12b expression at 36 hpf (S14 Fig.) suggesting that cxcl12b signals non-autonomously to cxcr4a expressing cells.

**Fig. 4:**
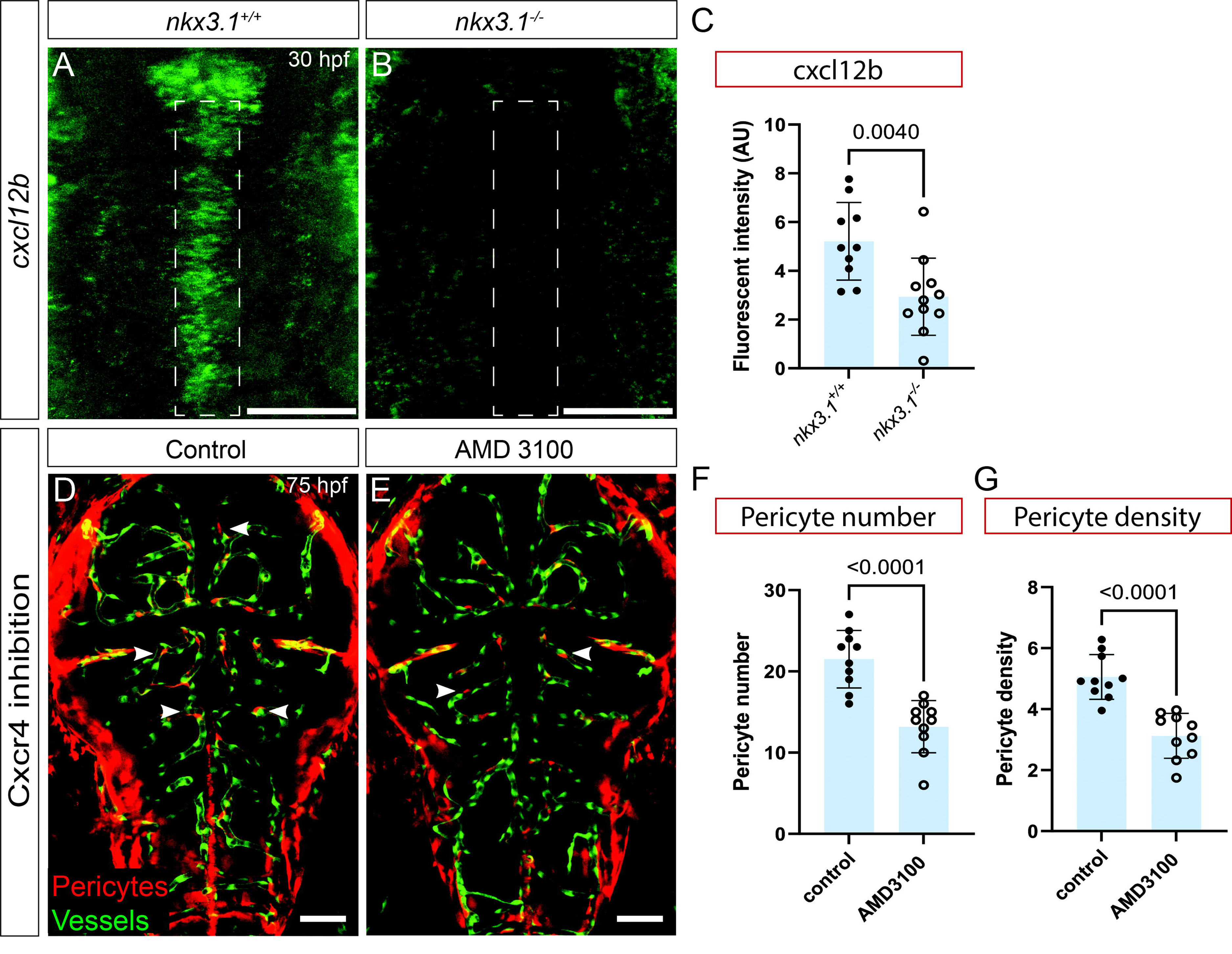
*cxcl12b* is regulated by *nkx3.1*; loss of Cxcr4 signaling reduces brain pericytes. All embryos were imaged dorsally in the head region. *cxcl12b* mRNA expression is reduced in the embryonic head of *nkx3.1^−/−^* mutants (B) as compared to wildtype controls (A) at 30 hpf as quantified by fluorescent intensity (C) of regions marked by dotted lines in A and B. (n=10 wildtype and 10 mutant embryos). Inhibition of Cxcr4 from 24-75 hpf using AMD 3100 leads to reduced brain pericytes in treated (E) vs. untreated (D) embryos (at 75 hpf (n=10). Pericytes (red cells, white arrows) are labeled by *TgBAC(pdgfr*β*:Gal4)*; *Tg(UAS:NTR-mCherry)*. Brain vessels in D and E are labeled by *Tg(flk:GFP; green)*. Quantification of (F) pericyte number and (G) pericyte density. (n=10 wildtype and 10 mutants). Statistical significance was calculated using the Student’s t-test. Scale bar is 50µm. The data underlying this figure can be found in S3 Table.

Supporting our results, single-cell sequencing of zebrafish at multiple stages as reported in the DanioCell database(32) shows expression of *nkx3.1, foxf2a* and *cxcl12b* at the 24-34 and 36-48 hpf at a time when *pdgfr*β is only weakly expressed (S15 Fig.). This independent dataset strongly supports the co-expression of these three genes in the pericyte lineage prior to definitive pericyte marker expression.

Taken together, our data shows that Cxcl12b is co-expressed with, and required downstream of Nkx3.1 in pericyte development.

### Cxcl12b function is required in *nkx3.1^+ve^* precursors for brain pericyte development

We next tested whether Cxcl12b is required in *nkx3.1^+ve^*precursors or Pdgfrβ^+ve^pericytes or in both cell types. We used transgenic overexpression of *cxcl12b* in *nkx3.1*^−/−^ mutants by constructing *Tg(UAS:cxcl12b;cryaa:mKate*) and crossing to Gal4 drivers in either the *nkx3.1^+ve^*precursor lineage or Pdgfrβ^+ve^ pericytes using *nkx3.1^−/−^;TgBAC(nkx3.1:Gal4)^ca101^*or *nkx3.1^−/−^;TgBAC(pdgfrb:Gal4FF)^ca42^*. We scored the number of brain pericytes in mutants carrying both transgenes. We find that *nkx3.1^−/−^* mutant pericyte numbers are increased by *nkx3.1*-driven *cxcl12b* overexpression (Fig 5A-C), but not by *pdgfr*β-driven *cxcl12b* overexpression at 75 hpf (Fig. 5E-G). Brain pericyte density is also increased when cxcl12b is expressed under the nkx3.1 driver (Fig 5D). Agreeing with the mRNA overexpression data, these experiments confirm the requirement for Cxcl12b in an early stage of pericyte differentiation in the *nkx3.1* pericyte precursor population but not in later pericyte development (*pdgfr*β).

**Fig. 5:**
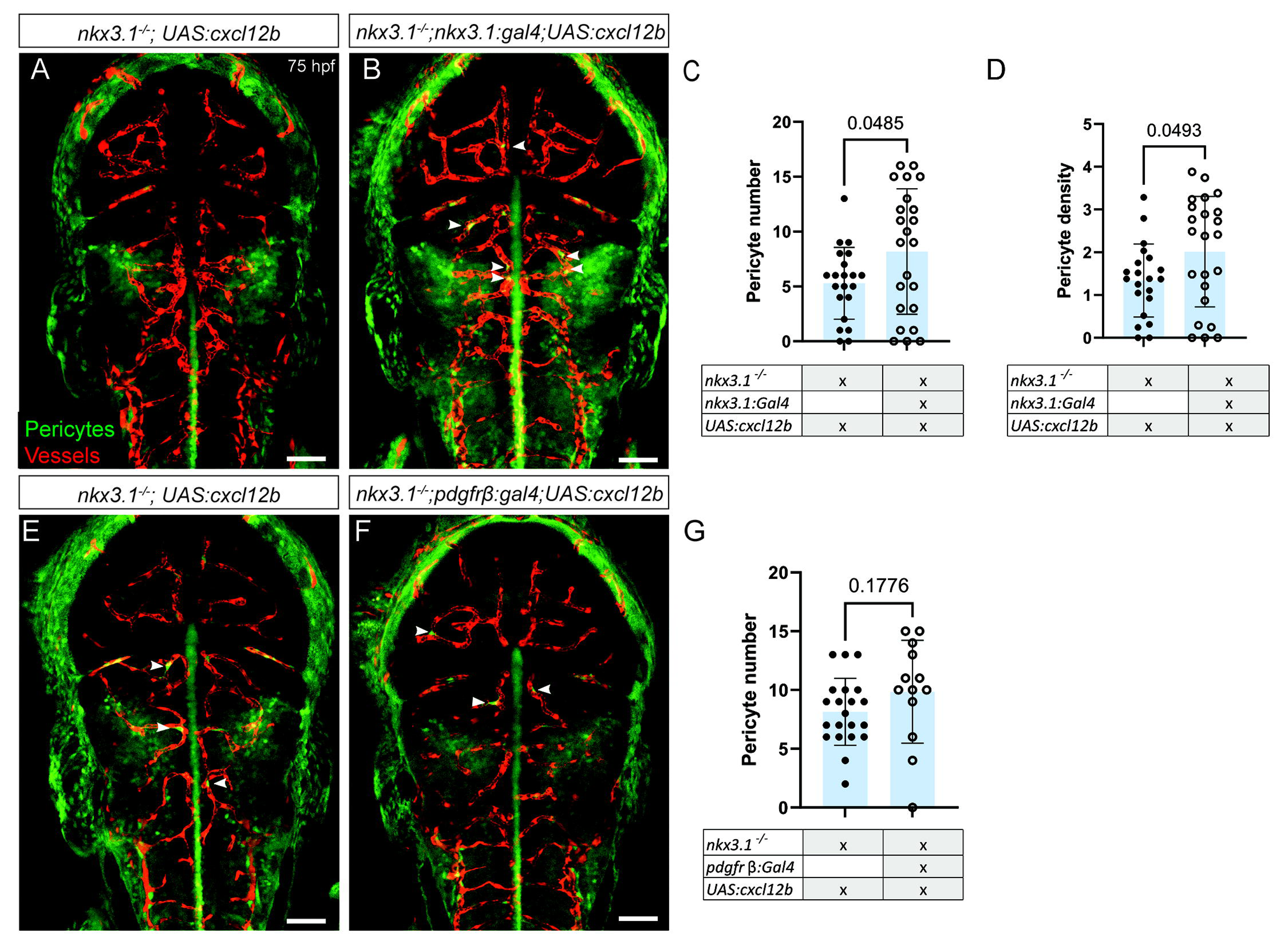
Re-expression of Cxcl12b increases pericyte numbers in *nkx3.1^−/−^*. All embryos were imaged dorsally in the head region. In comparison to a nkx3.1^−/−^ mutant without a Gal4 driver (A), expressing UAS:*cxcl12b* under the *nkx3.1* Gal4 driver *TgBAC(nkx3.1:Gal4)* increases pericyte numbers (B, C) and pericyte density (D) in *nkx3.1* mutants at 75 hpf (n=21 mutants without nkx3.1Gal4 and 22 with Gal4). In comparison to a nkx3.1^−/−^ mutant without a Gal4 driver (E), expressing UAS:*cxcl12b* under the under the *pdgfr*β Gal4 driver *TgBAC(pdgfr*β*:Gal4)* does not change pericyte numbers at 75 hpf (F,G; n=16 mutants without the pdgfrβ:Gal4, and 8 with pdgfrβ:Gal4). Arrowheads mark example brain pericytes (green) labeled by *TgBAC(pdgfr*β*:GFP)*. Brain vessels (red) are labeled by *Tg(kdrl:mCherry)*. Statistical significance was calculated using the Student’s t-test. Scale bar is 50µm. The data underlying this figure can be found in S3 Table.

**Fig. 6:**
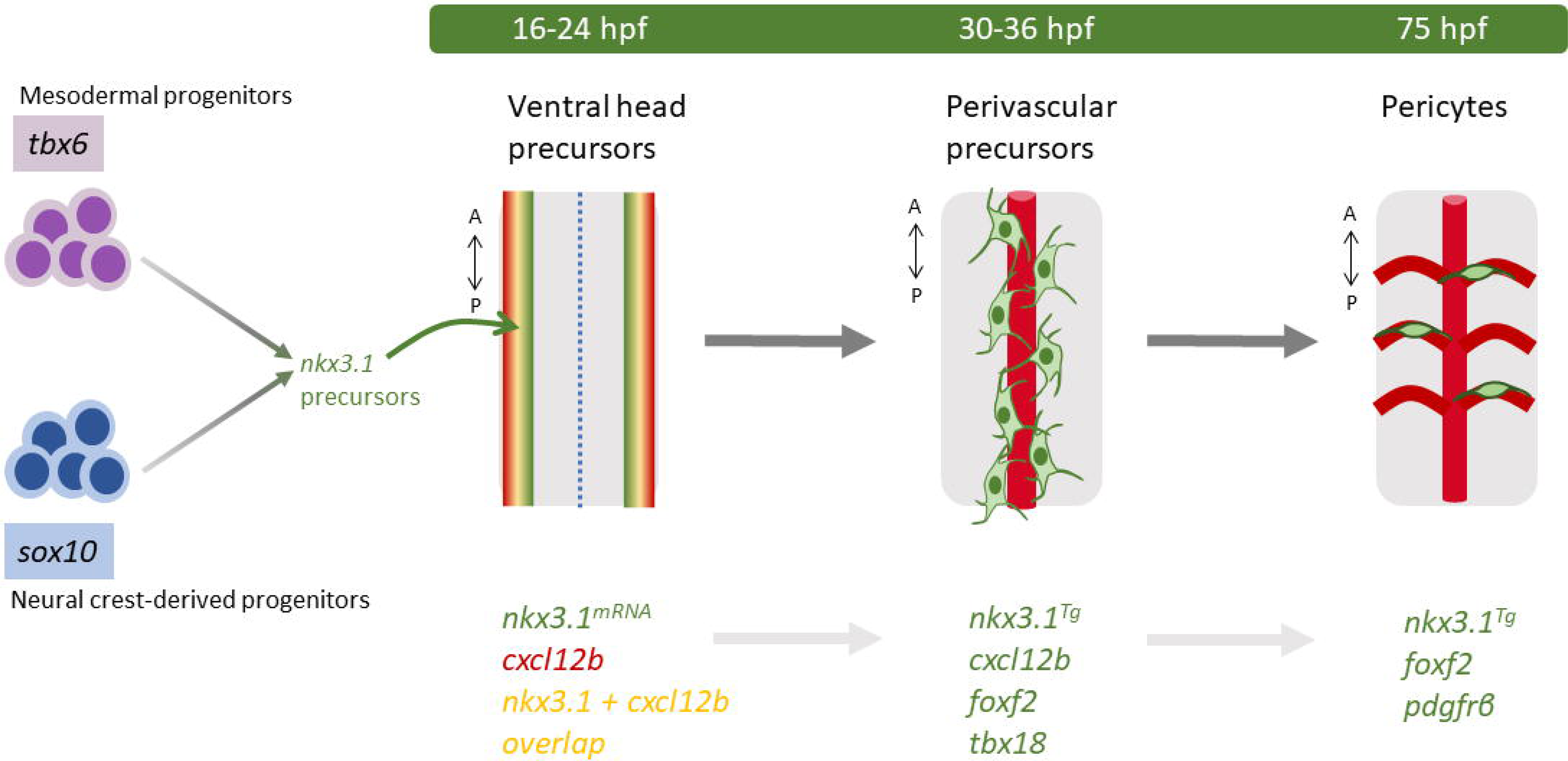
nkx3.1 is essential in pericyte precursors. Model of *nkx3.1* in pericyte differentiation. *nkx3.*1 is expressed in cells of both mesodermal and neural crest origin as determined by lineage analysis. All embryo schematics show a dorsal view of the hindbrain. At 16 and 24 hpf, nkx3*.1* and *cxcl12b* co-expressing precursors are located ventro-laterally. At 36 hpf, nkx3*.1*, *cxcl12b, tbx18 and foxf2* co-expressing precursors surround the developing basilar artery (BA). By 75 hpf pdgfrβ-expressing pericytes derived from *nkx3.1* precursors have migrated to the Central Arteries of the brain. Key genetic markers are indicated.

## Discussion

Little is known about the essential factors that contribute to the developmental journey of a differentiated pericyte. Most studies focus either on the upstream lineage origins from mesodermal or neural crest precursors, or on downstream genes important for differentiated pericytes but the intermediate genetic factors driving lineage differentiation are less well known. Here we show that *nkx3.1* is a vital intermediate gene in the differentiation journey of a pericyte. Using a transgenic reporter of *nkx3.1* expression we show that it precedes Pdgfrβ expression in the pericyte lineage. Cells in the brain that express *nkx3.1* become *pdgfr*β-expressing pericytes. Critically, *nkx3.1* is required for brain pericytes of both mesodermal and neural crest origin, suggesting that it is a gene that either unifies the convergent pericyte differentiation program from different lineages or is present in the newly unified population. Beyond the brain, previous work shows that *nkx3.1* is expressed in peri-vascular fibroblasts precursors of the trunk pericyte lineage (22). It is likely that *nkx3.1* is important for pericyte and fibroblast development in other areas of the embryo. Homozygous *nkx3.1* mutants are viable and fertile with reduced lifespan, suggesting that progeny of embryonic *nkx3.1*-expressing brain pericytes, trunk pericytes and/or fibroblasts likely contribute to progressive post-embryonic phenotypes. We show that the absence of *nkx3.1*-expressing cells results in a severe reduction in pericyte number. Loss of *nkx3.1* and gain of *nkx3.1* show that *nkx3.1* is both necessary and sufficient for modulating brain pericyte number. RNAseq and scRNAseq reveal the transcriptome of *nkx3.1*-expressing pericyte precursors as similar to fibroblasts and identify the chemokine cxcl12 as a critical factor for brain pericyte differentiation, whose expression is controlled by *nkx3.1*.

Brain pericytes are unusual as they arise from two distinct lineages, mesoderm and neural crest in both fish and mouse. No functional differences have been noted between pericytes from different origins. Using lineage tracing, we show that pericytes derived from both paraxial mesoderm (*tbx6*) and neural crest *(sox10*) express *nkx3.1,* and that pericytes derived from both mesoderm and neural crest are present in both the midbrain and hindbrain. Previous work has suggested a more compartmentalized contribution in fish where mesoderm contributes to hindbrain pericytes and neural crest contributes to midbrain pericytes (17). This may have occurred due to incomplete sampling in these difficult experiments, as similar reagents were used. The rarity of recombination events limits precise quantification of mesodermal and neural crest contribution, but contributions from both mesodermal (myeloid) and neural crest lineages to brain pericytes is also observed in mice (40).

Ablation of *nkx3.1* expressing cells or loss of the *nkx3.1* gene within these cells leads to an identical phenotype. Loss of *nkx3.1*-expressing cells or of *nkx3.1* leads to phenotypes associated with pericyte disruption including brain hemorrhage and reduced pericyte number (20, 26, 41), suggesting a critical role of Nkx3.1 in brain pericyte development. The level of pericyte loss is similar to that reported for both *foxf2* and *notch3* knockout fish suggesting that all three genes key players in pericyte differentiation (13, 20), potentially with some redundancy as *pdgfr*β knockout zebrafish have no pericytes, while all the models of transcription factor loss (nkx3.1, foxf2a, foxf2b) show reduced pericytes. Notch 3 gain of function increases pericyte numbers, and we show here that gain of Nkx3.1 leads to an increase in pericyte number. However, there was no change in notch3 expression in our RNA sequencing to indicate that there is a regulatory relationship, despite similar functions. Instead, Nkx3.1 and Notch3 may potentially act using similar downstream mechanisms (13).

Where and when *nkx3.1* acts in the pericyte differentiation cascade is an important question. We note that *nkx3.1* is an early embryonic gene and by 48 hpf its mRNA is undetectable, however perdurance of the *nkx3.1^NTR-mcherry^*transgenic allowed us to follow cells after the endogenous gene has turned off. Nothing is known of the Nkx3.1 protein and how long it may remain in an embryo, however, our data point to a specific early role in development. Incidentally, the first brain pericytes attach to the basilar artery by 36 hpf, and *nkx3.1* is expressed before this time.

To define the gene signature of this pericyte precursor population before pericyte markers are observed, we used single cell sequencing of sorted *nkx3.1^NTR-mCherry^* cells at 30 hpf. Analysis of single cells revealed *nkx3.1* contribution in thirteen gene clusters, of which the majority of *nkx3.1* positive cells belong to two precursor clusters and three fibroblast-like clusters as defined by expression of pan-fibroblast markers such as *col1a1, col5a1* and *pdgfr*α (Fig S7) (31). Interestingly, the Fb-V cluster is enriched in genes expressed in pericytes and/or crucial for their development, i.e., *tbx18* (35, 42), *cxcl12b* (37–39), and *foxf2a* (19–21). However, the Fb-V cluster lacks expression of ‘classical’ differentiated pericyte markers, including *Pdgfr*β *(17)*, *abcc9 (12, 43)*, *Kcnj8 (43)*, *ndufa4l2a and kcne4 (44)*. This indicates that Fb-V are fibroblast-like precursors and not differentiated cells. Differentiated pericytes are observed two days later in development. Of the two precursor clusters, Prog1 has extremely high expression of ribosomal protein subunits (30 *rps* (small ribosome) and 44 *rpl* (large ribosome) genes). Expression of *rps* and *rpl* genes is very enriched in mural cells from early human development (GW15-18) in comparison to later development (GW20-23) (28), suggesting that this cluster represents early precursors. The second precursor cluster, Prog2, overlaps in expression profile with all three fibroblast clusters (A, B, V). RNA velocity analysis suggest that Prog1 contributes to Fb-V and *nkx3.1*-expressing derivatives in the head, while Prog2 contributes to Fb-V, the two sclerotome fibroblast clusters Fb-A and Fb-B and heart mesoderm. RNA velocity also suggests that Fb-B is further differentiated from Fb-A. These two fibroblast clusters express classical markers of sclerotome, an expected major population of *nkx3.1*-expressing cells (22). Although it has a similar gene expression profile, sclerotome is spatially separate from brain pericytes in the embryo and forms the trunk mesenchyme. Extensive characterization of the pericyte transcriptome by scRNAseq in the adult mouse, differentiated fish pericyte (5 dpf), and embryonic human are all from later developmental stages than the transcriptome that we determine here, and therefore our analysis represents an early snapshot of pericyte differentiation (28, 44, 45).

To understand functional targets of Nkx3.1 in pericyte differentiation, we undertook bulk RNA sequencing of *nkx3.1* mutants. Among the downregulated genes, we found *ndufa4l2a* (a pericyte marker) (44), and *cxcl12b* are both downregulated in *nkx3.1^−/−^* embryos at 30 hpf. This is intriguing as single cell RNA sequencing showed expression of chemokine *cxcl12b* in *nkx3.1*-expressing fibroblasts, including Fb-V. Additionally, Cxcl12-cxcr4 signalling is involved in bone marrow-derived pericyte differentiation (46) and Cxcl12 also plays a role in recruitment of vascular smooth muscle cells to the zebrafish aorta (47), suggesting it is a strong candidate for mediating effects downstream of nkx3.1. *nkx3.1* positive pericyte precursors co-express *cxcl12b*, *tbx18, and foxf2a,* confirming our single cell RNA sequencing data. Functional small molecule inhibition of Cxcl12-Cxcr4 signaling significantly reduced pericyte numbers, and expressing *cxcl12b* either under promoters for *nkx3.1* (early precursors) or *pdgfrb* (more mature pericytes) showed that it was only able to increase pericyte numbers in *nkx3.1* mutants when expressed early (*nkx3.1* driver), but not once pericytes had differentiated to express (*pdgfrb* driver). Taken together, this suggests that expression of *nkx3.1* in a pericyte precursor promotes the expression of the chemokine *cxcl12* and influences pericyte differentiation. We propose a model where Cxcl12 released by pericyte precursors binds to Cxcr4a expressed by the basilar artery. Based on the model for smooth muscle cell recruitment (47), endothelial cells in turn might produce the Pdgfb ligand to facilitate attachment of Pdgfrβ+ pericytes on the basilar artery (48). Previous work suggesting that Pdgfb is attenuated with Cxcl12-Cxcr4 signalling inhibition supports our hypothesis (46).

Our study identifies a new player in pericyte differentiation, the transcription factor, *nkx3.1*. *nkx3.1* is required in an intermediate precursor cell state that exists temporally downstream of germ layer (mesoderm or neural crest) specification, and upstream of differentiated pericytes expressing canonical makers. The role of *nkx3.1* in brain pericyte precursors is transient and occurs in parallel to its role in trunk sclerotome, although these are distinct embryonic populations. We identify the key role of nkx3.1 in promoting the proper number of pericytes to emerge on brain vessels to promote downstream vascular stability. We show that expression of *nkx3.1* is necessary and sufficient to modulate developmental pericyte number, although it does not result in complete loss of pericytes, suggesting partial redundancy. Defining the novel gene expression signature of *nkx3.1*-expressing pericyte precursors using scRNAseq opens new avenues for understanding pericyte differentiation. For instance, expression of two transcription factors in the Fb-V cluster (*foxf2a, tbx18*) are associated with pericytes in previous studies (20, 21, 34, 35), although have poorly described roles, and regulatory changes in FOXF2 are associated with stroke in humans (20, 49). Future work focusing on identifying additional intermediate genes in pericyte differentiation is needed to illuminate the stepwise differentiation of pericytes from upstream precursors, and potential for regeneration in disease where pericytes are lost.

## Data availability

Bulk and single-cell RNA-Sequencing data reported in this work are available through NCBI GEO GSE232763.

## Methods

### Zebrafish

All procedures were conducted in compliance with the Canadian Council on Animal Care, and ethical approval was granted by the University of Calgary Animal Care Committee (AC21-0189). All experiments included wildtype or vehicle-treated controls as a comparison group and developmental stages, n’s, genotypes, and statistical outcomes are noted for each experiment. Embryos were maintained in E3 medium at 28C. For heat shock experiments, embryos were heated for 1 hour at 39C in a heating block.

The following published strains were used: *TgBAC(nkx3.1:Gal4)^ca101^ (23)*, *Tg(UAS:NTR-mCherry)^c264^* (50), *Tg(kdrl:GFP)^la116^* (51), *TgBAC(pdgfr*β*:GFP)^ca41^* (3), *TgBAC(pdgfr*β*:Gal4)^ca42^* (3), *Tg(kdrl:mCherry)^ci5^* (52).

The following strains were generated for this manuscript: *Tg(hsp70l:tBFP)^ca90^*, *Tg(hsp70l:tBFP-2a-nkx3.1)^ca91^, Tg(UAS:cxcl12a)^ca504^*. First, middle entry clones of tagBFP (Evrogen), tagBFP PCR-fused to *nkx3.1* or *cxcl12b* were made by amplification using primers in S4 Table and cloned into pDONR221 using BP Clonase (ThermoFisher). Transposon Tol2 vectors were assembled using LR Clonase (ThermoFisher) and the Tol2 vector (53). *Tg(tbx6:cre;myl7:GFP)^ca92^*and *Tg(sox10:cre;myl7:GFP)^ca93^* were injected in our laboratory from Tol2 plasmids provided by Tom Carney (54). *Tg(*hsp70l-loxP-STOP-loxP-H2B-GFP_cryaa-cerulean)^ca94^ was created from Addgene plasmid 24334 (55) by digestion with XhoI and re-ligation. This deleted mCherry but maintains the stop sequences and was followed by injection into zebrafish and raising of F1 and further generations from the F0 founder.

The *nkx3.1*^ca116^ mutant was generated using CRISPR/Cas9 as previously described(56). Target sites were identified using CHOPCHOP (57). The guide sequence was 5’-GGGGAGGCGGGAAAAAGAAGCGG −3’. To assemble DNA templates for sgRNA transcription, gene-specific oligonucleotides containing the T7 promoter sequence (5’-TAATACGACTCACTATA-3’), the 20-base target site, and a complementary sequence were annealed to a constant oligonucleotide encoding the reverse-complement of the tracrRNA tail. sgRNAs were generated by in vitro transcription using the Megascript kit (Ambion). Cas9 mRNA was transcribed from linearized pCS2-Cas9 plasmid using the mMachine SP6 kit (Ambion). To generate mutants, one-cell stage wild-type embryos were injected with a mix containing 20 ng/μl sgRNA and 200 ng/μl Cas9 mRNA. Injected fish were raised to adulthood and crossed to generate F1 embryos. T7 Endonuclease I assay (NEB) was then used to identify the presence of indel mutations in the targeted region of F1 fish. *nkx3.1*^ca116^ has a 13 bp deletion (S2 Fig.).

### Microscopy and imaging analysis

Embryos were sampled randomly from a clutch and included hemorrhaged and non-hemorrhaged embryos. Embryos were imaged with a 20x (NA 0.8) objective on an inverted Zeiss LSM880 Airyscan or LSM900 laser confocal microscope mounted in 0.8% low-melt agarose on glass-bottomed Petri dishes (MatTek). The imaging field was adjusted and tiled to encompass the entire brain vasculature (rostral-caudal and dorsal ventral) as marked with *Tg(kdrl:mCherry)*). For analysis, images were assembled using ImageJ (58). Briefly to identify pericytes without overlap, the stack was divided into two dorsal and ventral substacks that cover the entire depth of mid- and hindbrain CtAs (Central Arteries). Pericytes (defined as transgene-labelled cells with a soma and processes) were counted manually on these stacks ensuring that no pericyte was counted twice. The numbers were compiled to obtain the total number of pericytes. Blood vessel network length was calculated automatically using VesselMetrics, a Python program (59). All computational analysis was accompanied by manual inspection of the data, removing any erroneous segments (unlumenized angiogenic sprouts). Pericyte density was obtained by dividing the number of pericytes over the vessel network length.

### Statistical analysis

Statistical analysis used GraphPad Prism 7 software, using a one-way ANOVA with multiple comparisons and a Tukey’s or Dunnett’s post-hoc test. Results are expressed as mean ± SD. The N’s of biological and n’s of technical replicates, p-values and statistical test used are provided in the figure legends. All raw data underlying figures can be found in S3 Table.

### Lineage analysis

Stable transgenic reporter lines harboring *Tg(*hsp70l-loxP-STOP-loxP-H2B-GFP_cryaa-cerulean)^ca94^ (nuclear, GFP), (TgBAC(pdgfrb:Gal4FF) or TgBAC(nkx3.1:Gal4)) and Tg(UAS:ntr:mCherry) (cytoplasmic, pericyte, red) were crossed with *Tg(tbx6:cre;myl7:GFP)^ca92^* or *Tg(sox10:cre;myl7:GFP)^ca93^*. Fish were imaged at 75 hpf and double positive pericytes counted separately in midbrain and hindbrain.

### In situ hybridization

For Hybridization Chain Reaction (HCR) in situ hybridization, custom probes for *nkx3.1, tbx18, foxf2a*, *cxcr4a* and *cxcl12b* were obtained from Molecular Instruments (Los Angeles, CA). The in-situ staining reactions occurred as recommended by the manufacturer.

### Drug treatments

All small molecules are listed in S5 Table. AMD 3100 was dissolved in E3 zebrafish water (5 mM NaCl, 0.17 mM KCl, 0.33 mM CaCl_2_, 0.33 mM MgSO_4_, 0.00001 % Methylene Blue) and applied at 100 µM from 24-75 hpf in the dark. Metronidazole was dissolved at 5mM n E3 was applied from 24-48 hpf in the dark. Embryos were washed out and grown in E3 medium without the drugs until imaging.

### Embryo dissociation and FACs sorting

Two biologically independent replicates were dissociated, FACs sorted, library prepped and sequenced independently. Each replicate started with ∼250 wildtype 30 hpf zebrafish embryos on the TL background expressing the *Tg(nkx3.1:Gal4;UAS:NTR-mCherry)^ca101^*transgene (22). Embryos were anaesthetized in Tricaine, kept on ice, and then dissociated with 0.25% Trypsin in 1mM EDTA at 28C with 300 rpm shaking for 20 minutes. The reaction was stopped using 1.25 µL 0.8M CaCl_2_ and 100 µL 1% FBS. Cells were washed in 1%FBS in Dulbecco’s PBS 3 times before straining through 75 µm and 30 µm strainers (Greiner Bio-One and Miltenyi Biotec) in Dulbecco’s PBS. Cell viability was determined using Trypan blue (Sigma) and was greater than 80% for both replicates. Cells were subjected to FACS (BD Facs Aria III, BD Biosciences) sorting for mCherry and excluding doublets and non-viable cells.

### scRNA-Seq library construction, sequencing

The biologically independent replicates of FACS-sorted nkx3.1-positive cells were each made into an independent library using the Chromium Single Cell Chip A kit, the Chromium Single cell 3′ Library & Gel beaded kit V3 and 3.1 Next GEM (10x Genomics; Pleasanton, CA, USA). Libraries were sequenced on the Illumina NovaSeq using an S2 Flowcell (Illumina) at the Centre for Health Genomics and Informatics, University of Calgary. A total of 3359 cells passed quality control for analysis, 2129 cells from sample 1 and 1230 from sample 2. The data from both samples was integrated using Cellranger_aggr. All raw FASTQs were aligned to the zebrafish reference genome GRCz11 (version 4.3.2) generated using the Cellranger version 3.1.0 mkref pipeline. The gene-barcode matrix output was processed using a standard Seurat v3 pipeline in R (60). Quality control steps filtered out cells expressing fewer than 200 genes or more than 2500 genes, as well as cells that had a mitochondrial content <5%. Uniform manifold approximation and projection (UMAP) with 15 principal components and resolution of 0.6 was used for dimension reduction. To assess the reproducibility of the two samples, cells from each replicate were projected onto the UMAP and proportions of cells in each cluster compared between samples (S6 Fig.). The R code for this analysis is in the Supplemental Data.

### Cluster assignment

Clusters were manually assigned by querying top markers in each cluster with known markers from annotated datasets (28-31, 61). A list of the cluster number, assignment, genes used for assignment and references for known cluster markers can be found in S1 Table.

### RNA velocity analysis

The RNA velocity pipeline for Seurat objects (Satija lab) was implemented. Loom files were generated from position-sorted aligned BAM files using velocyto (version-0.17.15) (62). A combined loom file containing counts of spliced, unspliced and ambiguous transcripts of both samples was read using Seurat function ReadVelocity. A count table of spliced and unspliced transcripts (stored as a Seurat object) was processed to merge with the existing Seurat object generated for cluster identification. Velocity analysis was performed on the integrated Seurat object using velocyto.R (62). RNA velocity vectors were projected on UMAP embeddings generated during cluster generation. The python code for this analysis is in Supplemental data.

### Bulk RNA sequencing

30 hpf MZ *nkx3.1^ca116^* mutant and wild type embryos were collected and RNA prepared using Trizol (ThermoFisher, Waltham MA). Libraries were prepared from 50ng/µg/µl of total mRNA per sample using the Illumina Ultra II directional RNA library prep (SanDiego, CA), and sequenced via the NovaSeq SP 100 cycle v 1.5 sequencing run with 33 million reads per sample. 4 replicates were sequenced per condition.

For bulk RNASeq, quality control evaluations performed by FastQC v0.11.9 (63) revealed that the reads were of high quality and required no trimming step. Consequently, the reads were mapped to the reference genome danRer11 obtained from UCSC and gene annotation file Zebrafish ENSEMBL Lawson v4.3.2 (64) using the splice-aware alignment tool, STAR v2.7.8a (65). STAR was run on default settings with the additional optional command ‘--quantMode GeneCounts’ to generate the gene counts files. The gene counts files were filtered for genes with less than 10 counts and normalized to the median ratio using DESeq2 v1.34.0 (66). Differentially expressed genes were identified as genes with FDR-adjusted p-value < 0.05 and log2(fold-change) < −0.5 or log2(foldchange) > 0.5. Volcano plots of differentially expressed genes was generated using Enhanced Volcano v1.12.0 (67).

**S1 Movie:** Timelapse of labelled Tg(*nkx3.1^NTR-^mcherry)* perivascular cells as they migrate and proliferate on blood vessels from 55-65 hpf.

**S1 Table.** 30 hpf scRNAseq (see Excel sheet)

**S2 Table.** 30 hpf bulk RNAseq (see Excel sheet)

**S3 Table:** Raw data (see Excel Spreadsheet)

**S4 Table:**
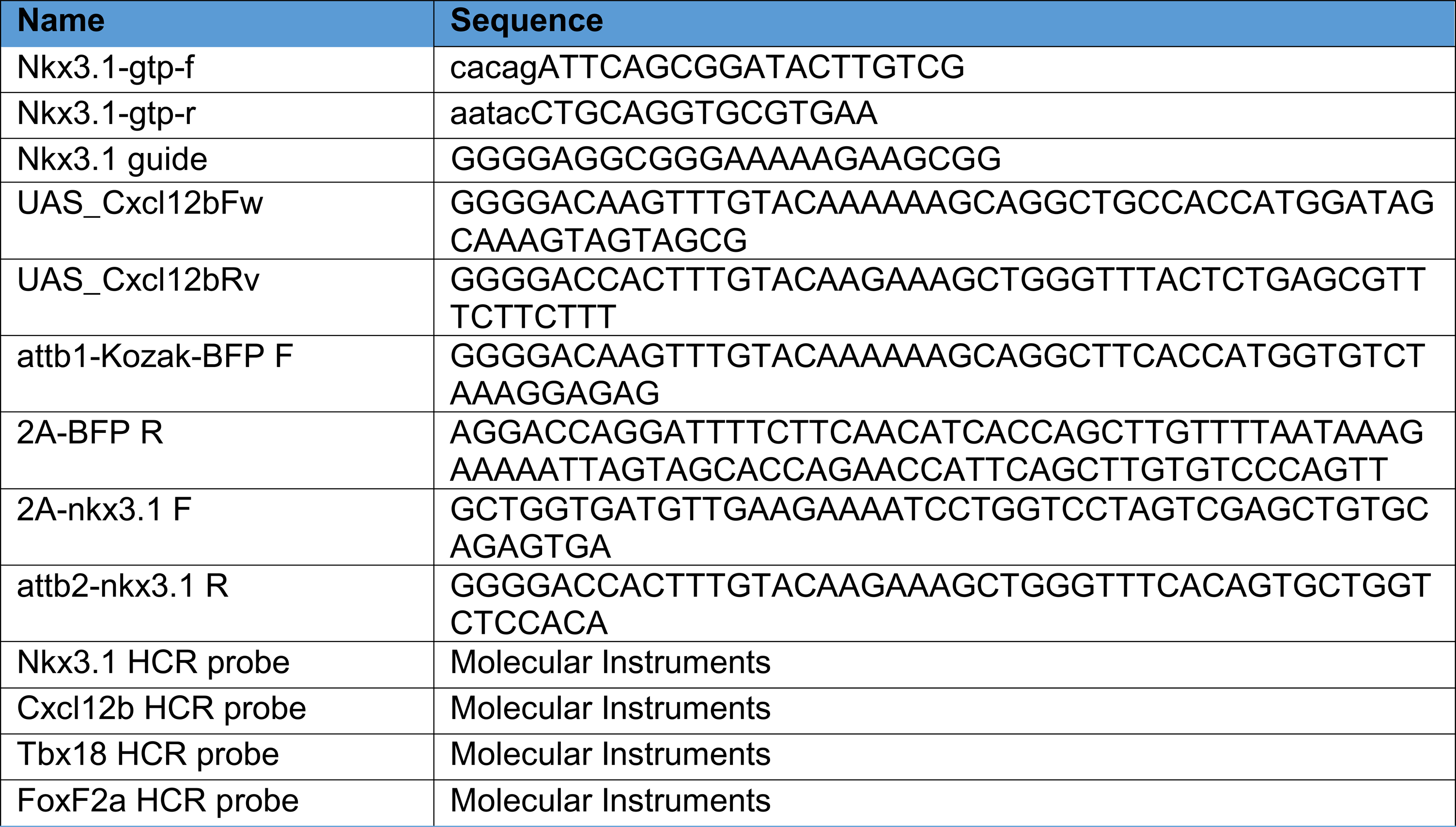
Primers, guides, HCR probes.

**S5 Table:** Reagent Table.

| Chemical name | Source | Catalog number | Concentration |
| --- | --- | --- | --- |
| Metronidazole | Sigma | M3761 | 5mM |
| AMD 3100 | Sigma | A5602 | 100 µM |
| Chromium Single Cell Chip A kit | 10X Genomics | 120236 |  |
| Chromium Next GEM Single Cell 3' GEM, Library & Gel Bead Kit v3.1 | 10X Genomics | 1000075/ 1000128 |  |

