## Supplemental figures for "The development of brain pericytes requires expression of the transcription factor *nkx3.1* in intermediate precursors"

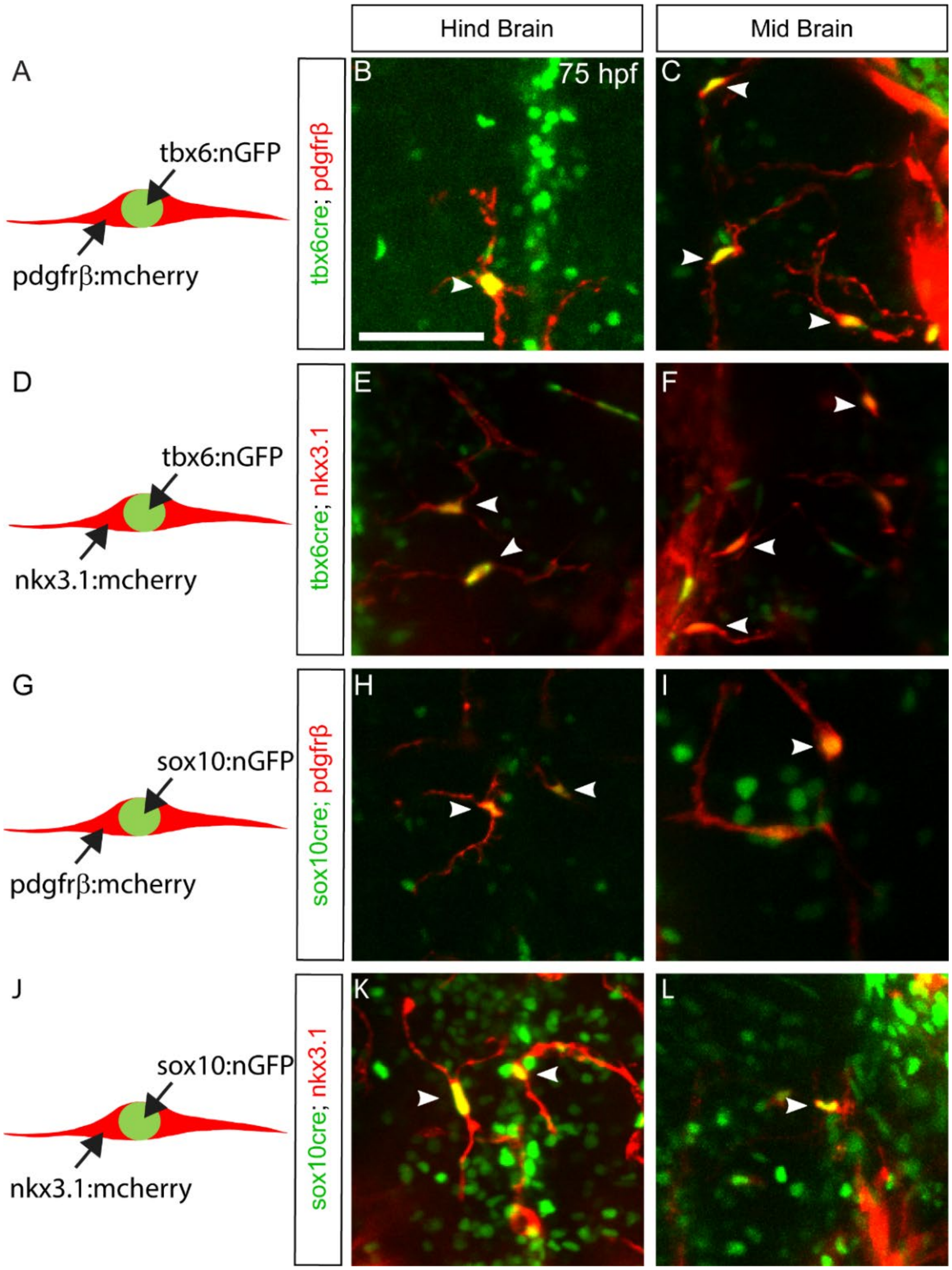


**Fig. S1: Lineage tracing of nkx3.1-expressing cells and pericytes at 75 hpf.** (A, D, G, J) Schematics of lineage tracing strategy showing how Tg(*tbx6:Cre)* or Tg(*sox10:Cre*) drivers crossed to the *Tg(loxp-stop-loxp-H2B-GFP*) reporter labels progeny with nuclear GFP. Pericytes *TgBAC(pdgfrb:Gal4FF) or nkx3.1-expressing cells TgBAC(nkx3.1:Gal4)* + *Tg(UAS:ntr:mCherry)* label cytoplasm red. (B-L) All images are dorsal views of embryonic mid or hind brains at 75 hpf, as marked. (B, C, E, F) Mesodermal lineage trace of pericytes (B, C) and Nkx3.1 cells (E, F). (H, I, K, L) Neural crest lineage trace of pericytes (H,I) and Nkx3.1 cells (K, L). Arrowheads indicate double positive *pdgfrβ* or *nkx3.1* cells. Scale bar is 50μm.

**A. DNA sequence around mutation**

```
Nkx3.1 wildtype:   CGGGGGGAGGCGGGAAAAAGAAGCGGTCGCGCGCCGCGT
Nkx3.1ca116:       CGGGGGGAGGCGGTCGCGCGCCGCGT
```

**B. Protein sequence (teal= homeobox, grey = out of frame translation)**

```
Nkx3.1wt       MATSNKQLTSFFIEDILSLKEDKKDEDSCNAESDRDDSTDRQTDSADTCRTSEGKTVSST
Nkx3.1ca116    MATSNKQLTSFFIEDILSLKEDKKDEDSCNAESDRDDSTDRQTDSADTCRTSEGKTVSST

Nkx3.1wt       EMTGGGGKKKRSRAAFTHLQVLELEKKFSRQRYLSAPERTHLASALHLTETQVKIWFQNR
Nkx3.1ca116    EMTGGGGRAPRSRTCRFWSWRRSSAVSGT*

Nkx3.1wt       RYKTKRRQLTTEHSKDYFQKSNAAAMAATEEDFFRASLLATVYKSSPYRPYVYDLHGLSM

Nkx3.1wt       WRPAL*
```

**C. Schematic of protein domains**

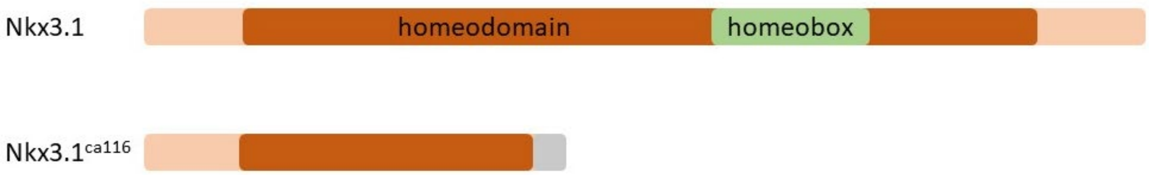


**Fig. S2: Nkx3.1$^{ca116}$ mutation characterization**

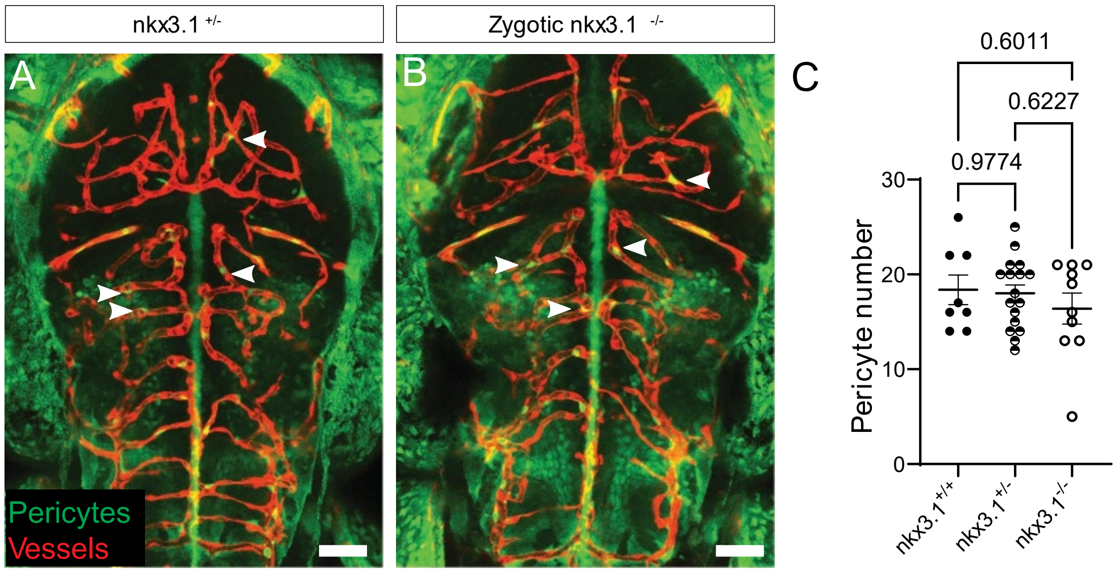


**Fig. S3: Zygotic *nkx3.1* mutants have wildtype pericyte numbers at 75 hpf.**

(A-B) Dorsal views of embryonic brain of zygotic *nkx3.1* mutants showing no change in brain pericyte numbers as compared to wildtype. Pericytes (green, arrowheads) are labelled with *TgBAC(pdgfrβ:GFP)* and vessels (red) are labelled with *Tg(kdrl:mCherry)*. (C) Quantitation of pericyte numbers shows no significance using one-way ANOVA with Tukey's test. (n=8 wildtypes, 17 heterozygotes and 10 mutants). Scale bar is 50 µm. The data underlying this figure can be found in Supp Table 3.

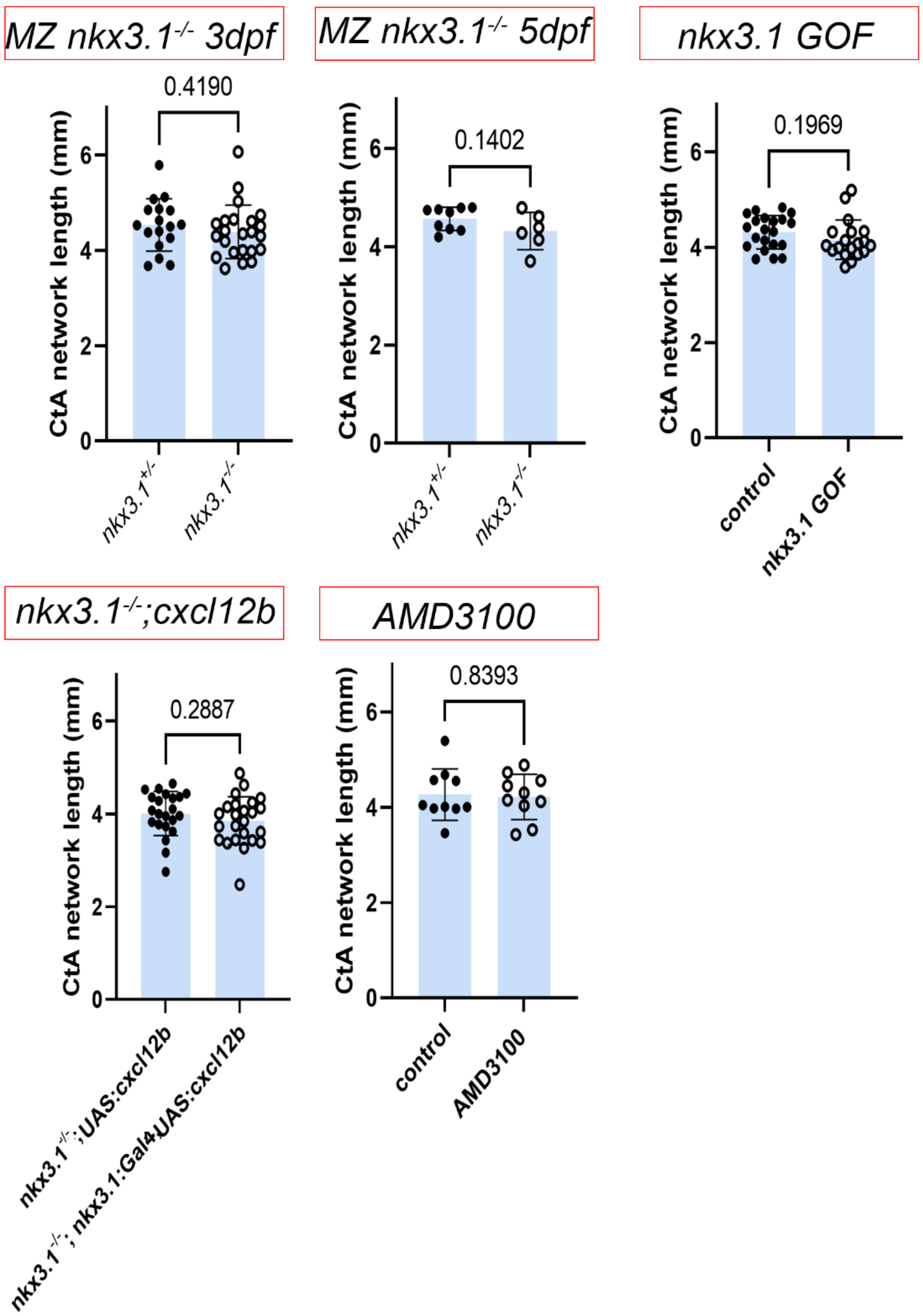


**Fig. S4: Central Artery vessel network length is unchanged across multiple experimental manipulations**

Vessel network length of the hindbrain CtAs was measured using VesselMetrics. Genotypes and treatments are labelled. No treatment or mutant significantly alters the endothelial vessel length. Statistics used a Student's t-test. The data underlying this figure can be found in Supp Table 3.

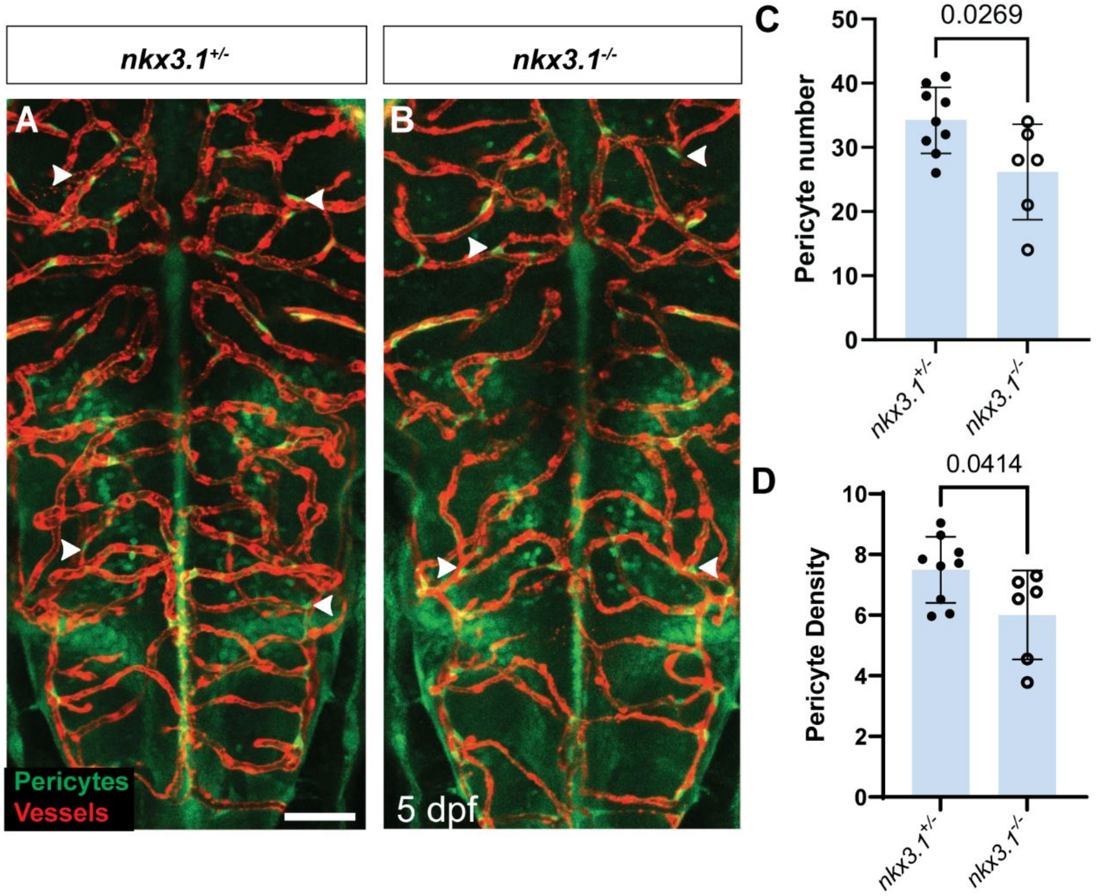


**Fig. S5: Pericyte number and density is decreased at 5dpf in *nkx3.1* maternal-zygotic mutants**

(A-B) Dorsal views of embryonic brain of MZ *nkx3.1* mutants at 5 dpf. Pericytes (green, arrowheads) are labelled with *TgBAC(pdgfrβ:GFP)* and vessels (red) are labelled with *Tg(kdrl:mCherry)*. There are significantly fewer brain pericytes (C) and reduced pericyte density (D) in nkx3.1$^{-/-}$ mutant as compared to nkx3.1$^{+/-}$ controls. Statistics used a Student's t-test. (n=9 wildtypes, 6 mutants). Scale bar is 50 µm. The data underlying this figure can be found in Supp Table 3.

A

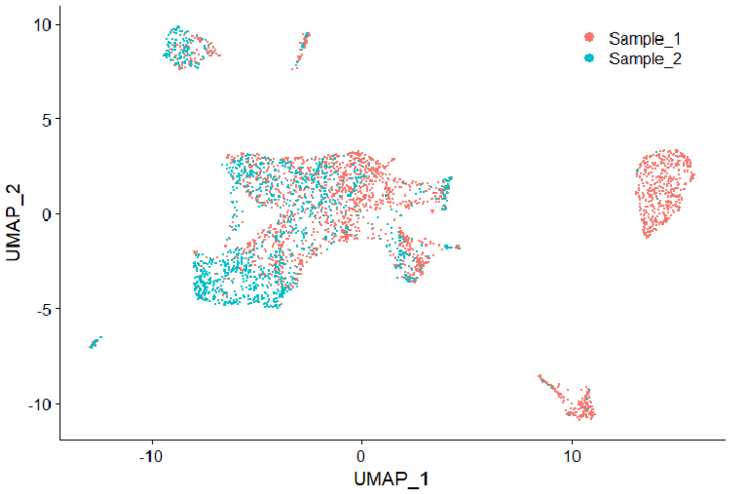


B

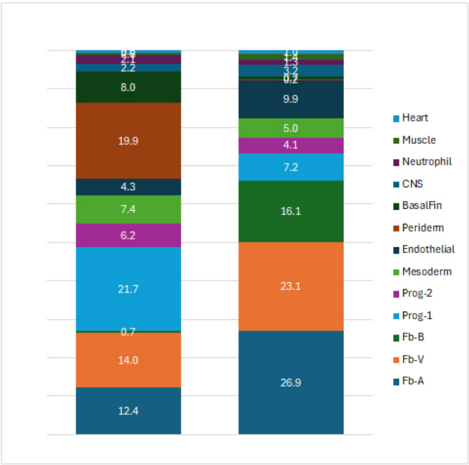


| Cluster | Sample_1 | Sample_2 |
|---|---|---|
| Fb-A | 12.4 | 26.9 |
| Fb-V | 14.0 | 23.1 |
| Fb-B | 0.7 | 16.1 |
| Prog-1 | 21.7 | 7.2 |
| Prog-2 | 6.2 | 4.1 |
| Mesoderm | 7.4 | 5.0 |
| Endothelial | 4.3 | 9.9 |
| Periderm | 19.9 | 0.2 |
| BasalFin | 8.0 | 0.7 |
| CNS | 2.2 | 3.2 |
| Neutrophil | 2.1 | 1.3 |
| Muscle | 0.6 | 1.4 |
| Heart | 0.6 | 1.0 |

**Fig. S6: Analysis of proportions and batch effects analysis of scRNAseq**

(A) Overlay of UMAP projections from two biologically independent scRNAseq samples (1, red, 2, blue) showing cells with similar distributions from both samples. (B) Stacked barplot showing the proportion of cells in each cluster deriving from the first sample (1) or second sample (2) as normalized as a percentage to the total number of cells from each sample. (Right) Table of proportions of the different cell types in each sample.

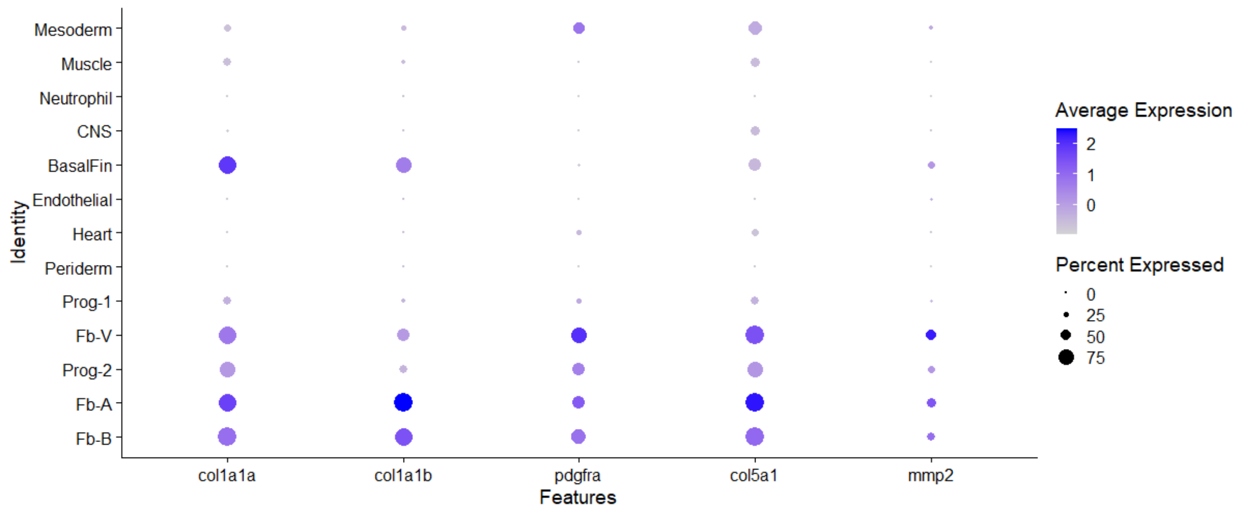


**Fig. S7: Dotplot showing expression of common fibroblast markers in Progenitor-2 and Fb clusters**

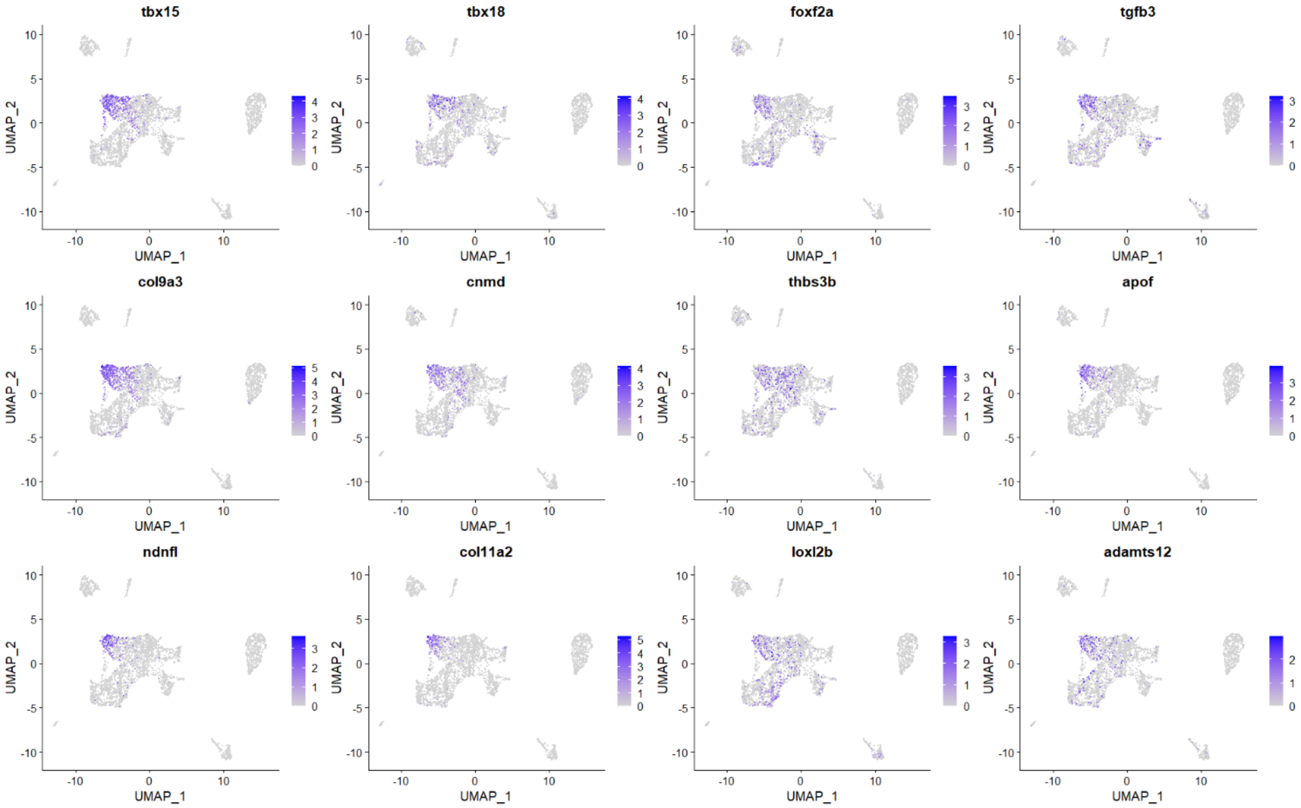


**Fig. S8: Featureplots of genes enriched in the Fb-V scRNAseq cluster (fibroblast-like pericyte precursors)**

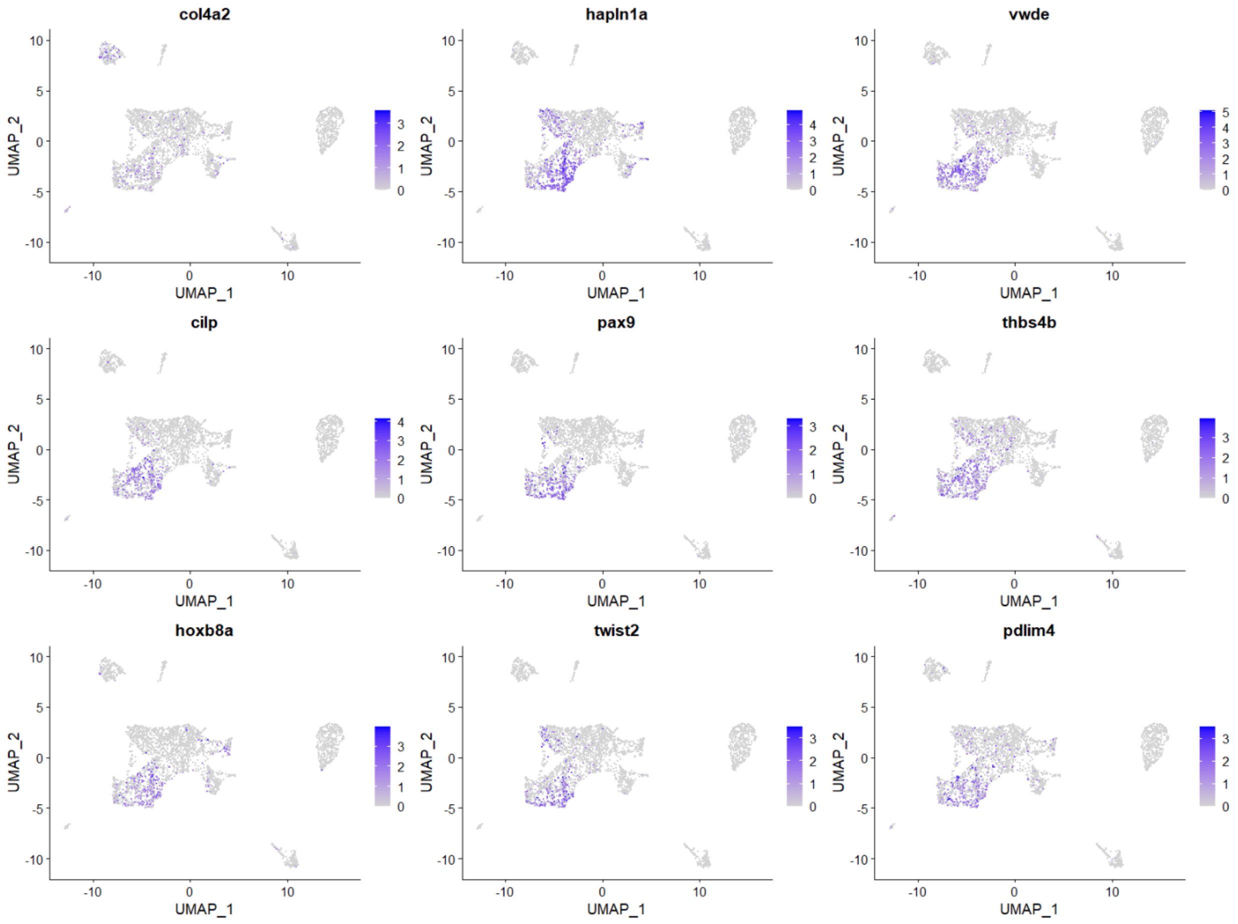


**Fig. S9: Featureplots of genes enriched in the Fb-A scRNAseq cluster**

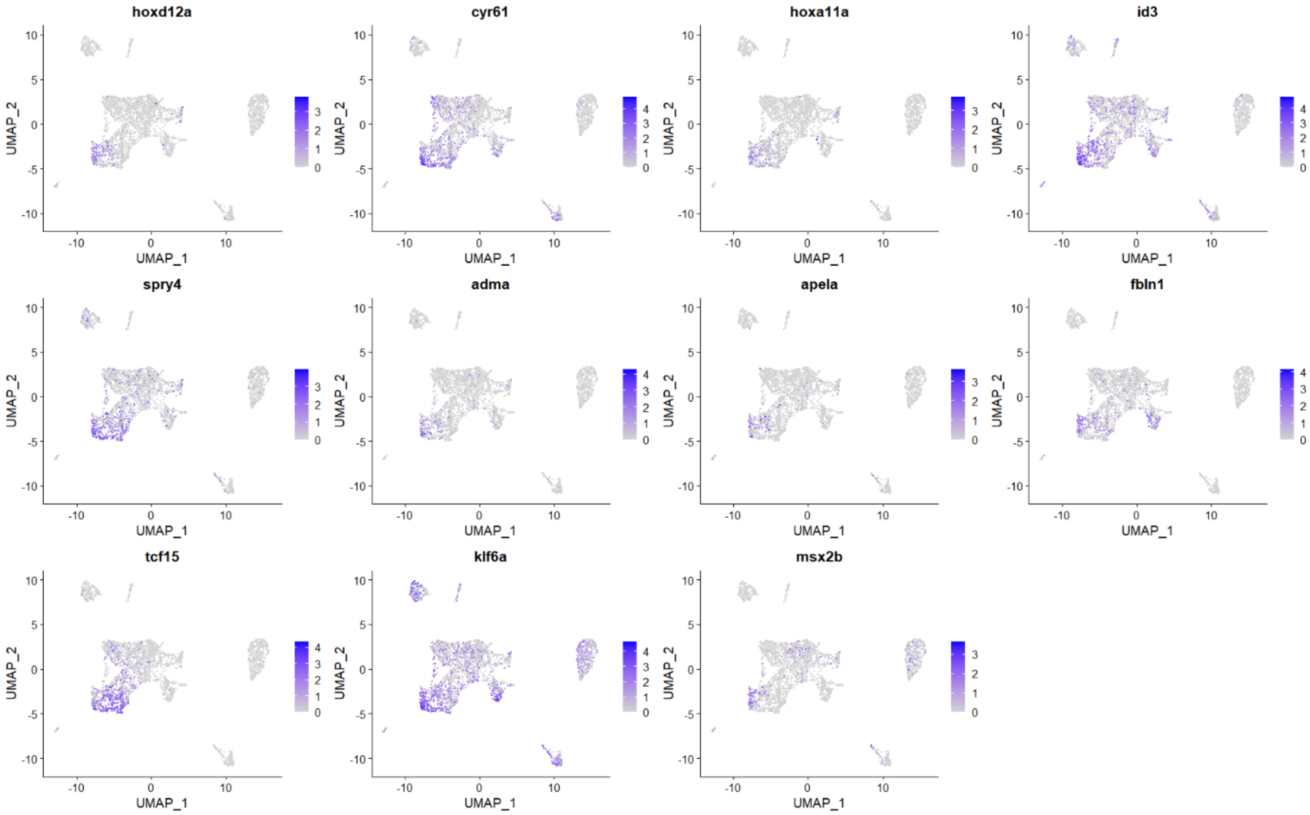


**Fig. S10: Featureplots of genes enriched in the Fb-B scRNAseq cluster**

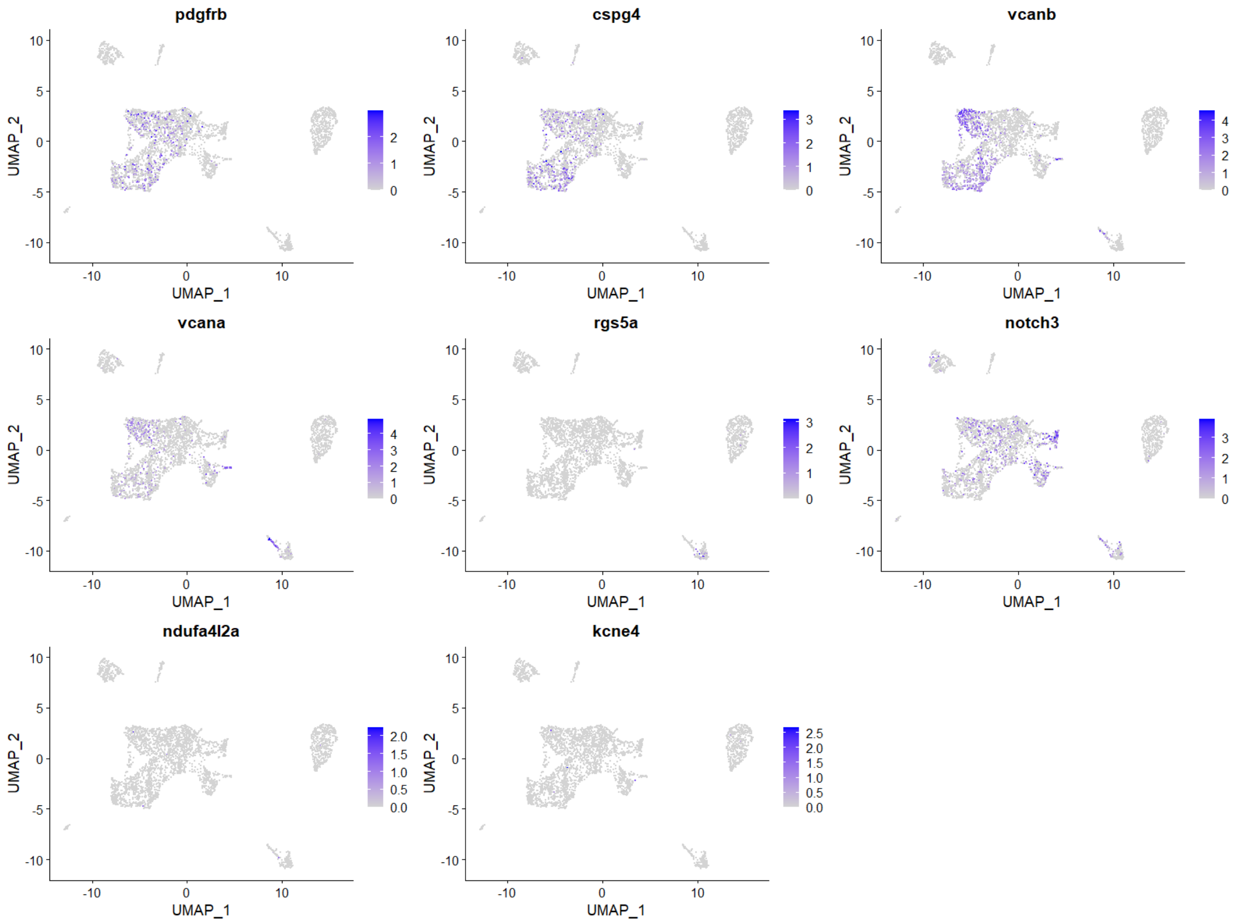


**Fig. S11: Featureplots of canonical pericyte markers in the nkx3.1-positive cell scRNAseq data.**

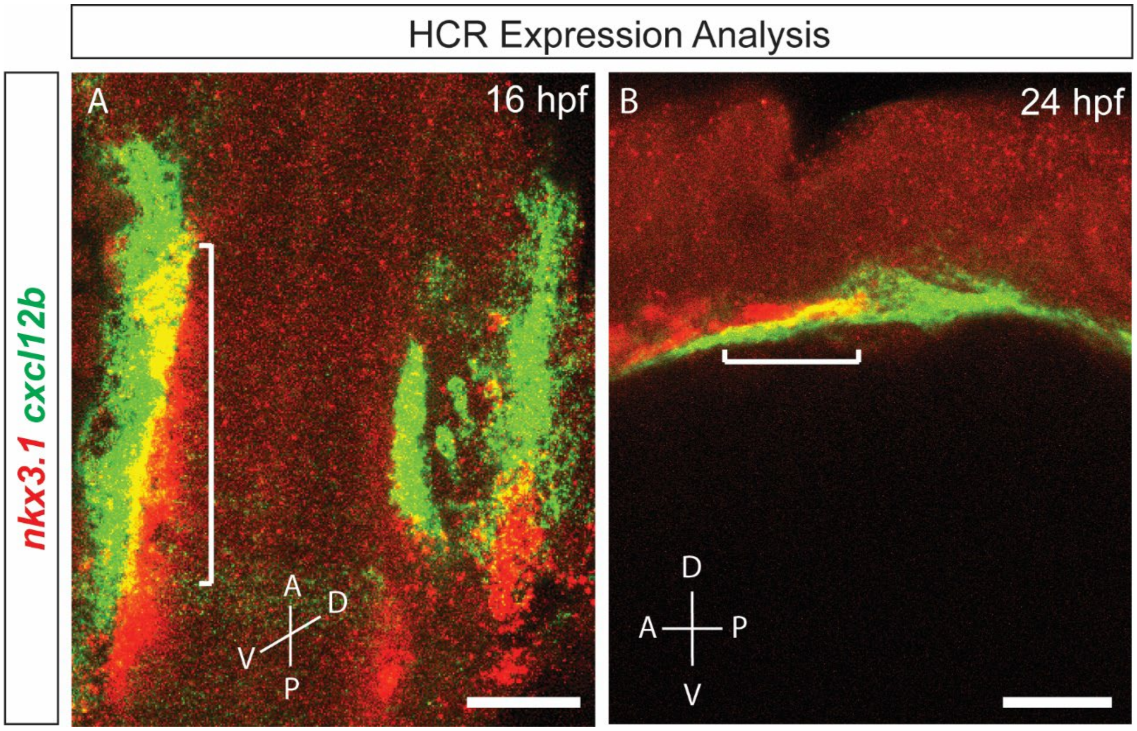


**Fig. S12: Expression analysis of *nkx3.1* and *cxcl12b* using HCR.** (A) Dorsal view of the posterior head region of 16 hpf embryo showing expression overlap (white bracket) between *cxcl12b* (green) and *nkx3.1* (red). (B) Lateral view of the posterior head region of 24 hpf embryo showing expression overlap (white bracket) between *cxcl12b* and *nkx3.1*. A-Anterior, P-Posterior, D-Dorsal, V-Ventral. Scale bar is 50 μm.

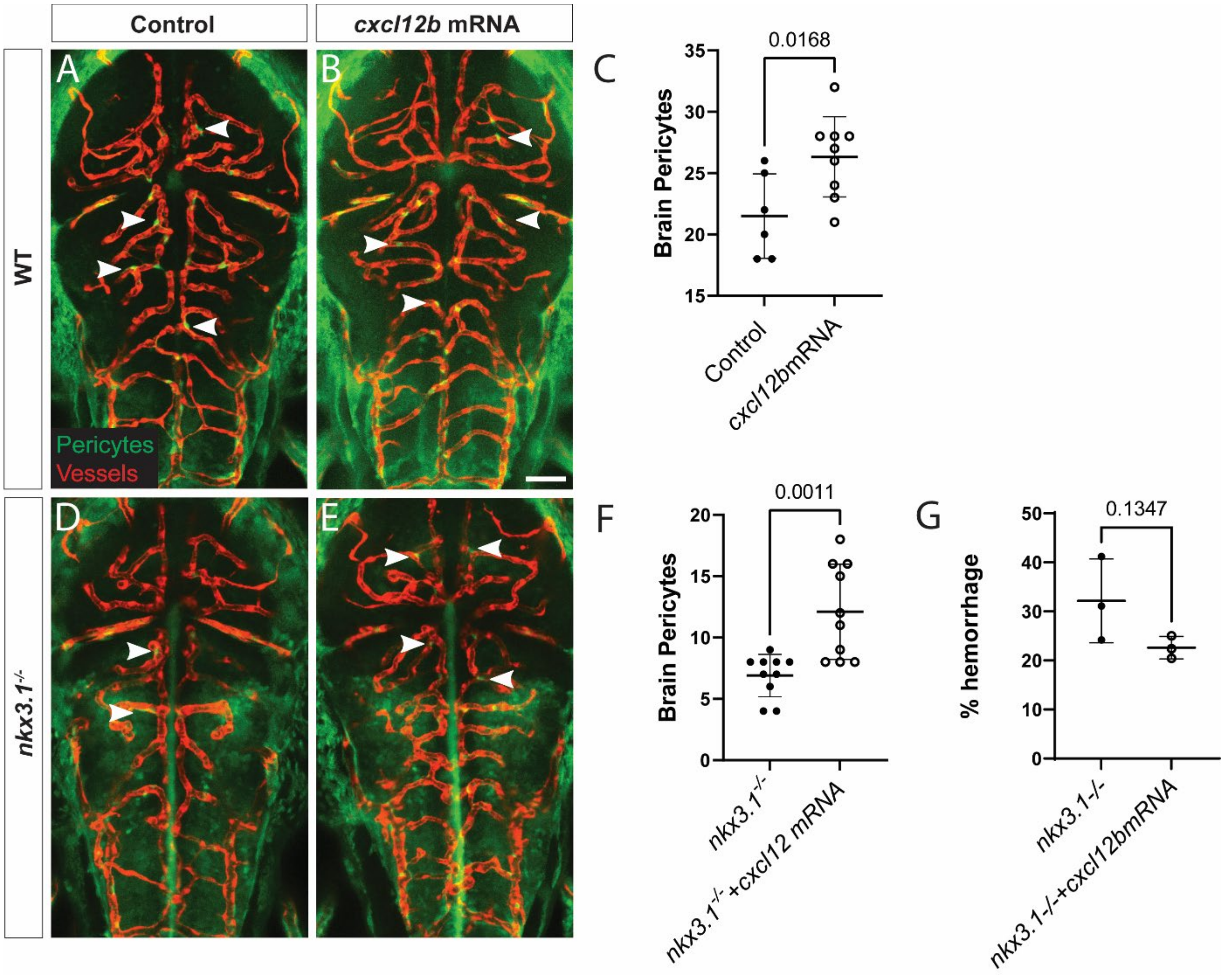


**Fig. S13: *cxcl12b* mRNA injection increases pericyte numbers at 75 hpf.** (A-C) Dorsal views of embryonic brain of uninjected control (A) and *cxcl12b* mRNA injected (B) embryos showing an increase in brain pericyte number (C) Quantitation of brain pericytes (n= 5 wildtype and 9 injected embryos). (D-F) Dorsal views of embryonic brain of uninjected *nkx3.1*$^{-/-}$ (D) and *cxcl12b* mRNA injected *nkx3.1*$^{-/-}$ embryos (E) showing an increase in brain pericyte number due to cxcl12b mRNA injection (F, n=10 uninjected and 10 injected mutants). Pericytes (green, arrowheads) are labelled with *TgBAC(pdgfrβ:GFP)* and vessels (red) are labelled with *Tg(kdrl:mCherry)*. (G) Hemorrhage rates were unchanged in mutants at 48 hpf (N=3 with 141 uninjected at 167 injected mutants analyzed). Statistics used the Students T-test. Scale bar is 50 μm. The data underlying this figure can be found in Supp Table 3.

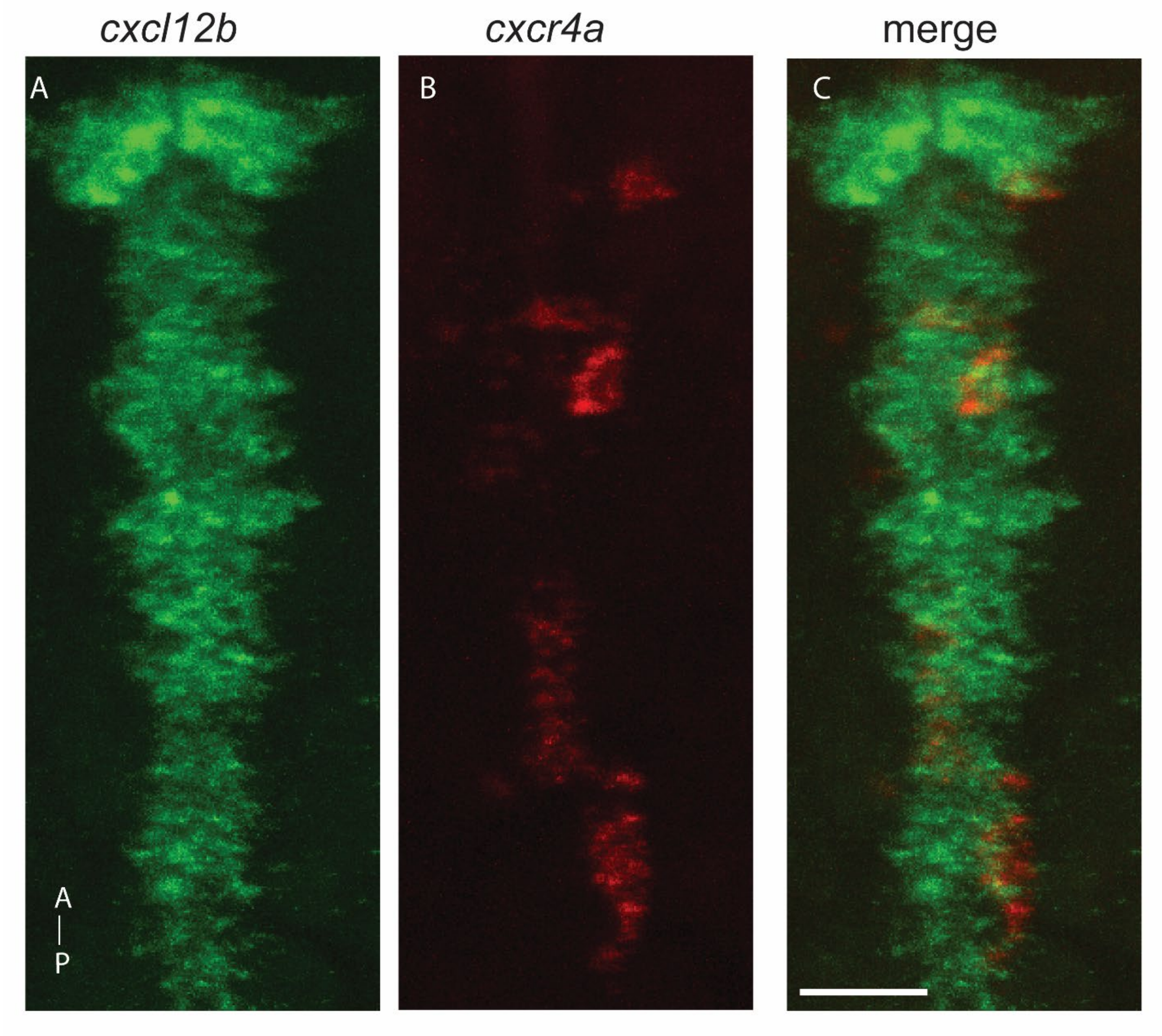


**Fig. S14: No overlap between *cxcl12b* and cxcr4a mRNA at 36 hpf**

Dorsal view of the ventral head region of 36 hpf embryo showing no expression overlap between *cxcl12b* (green) and *cxcr4a* (red). A-Anterior, P-Posterior, Scale bar is 20 μm.

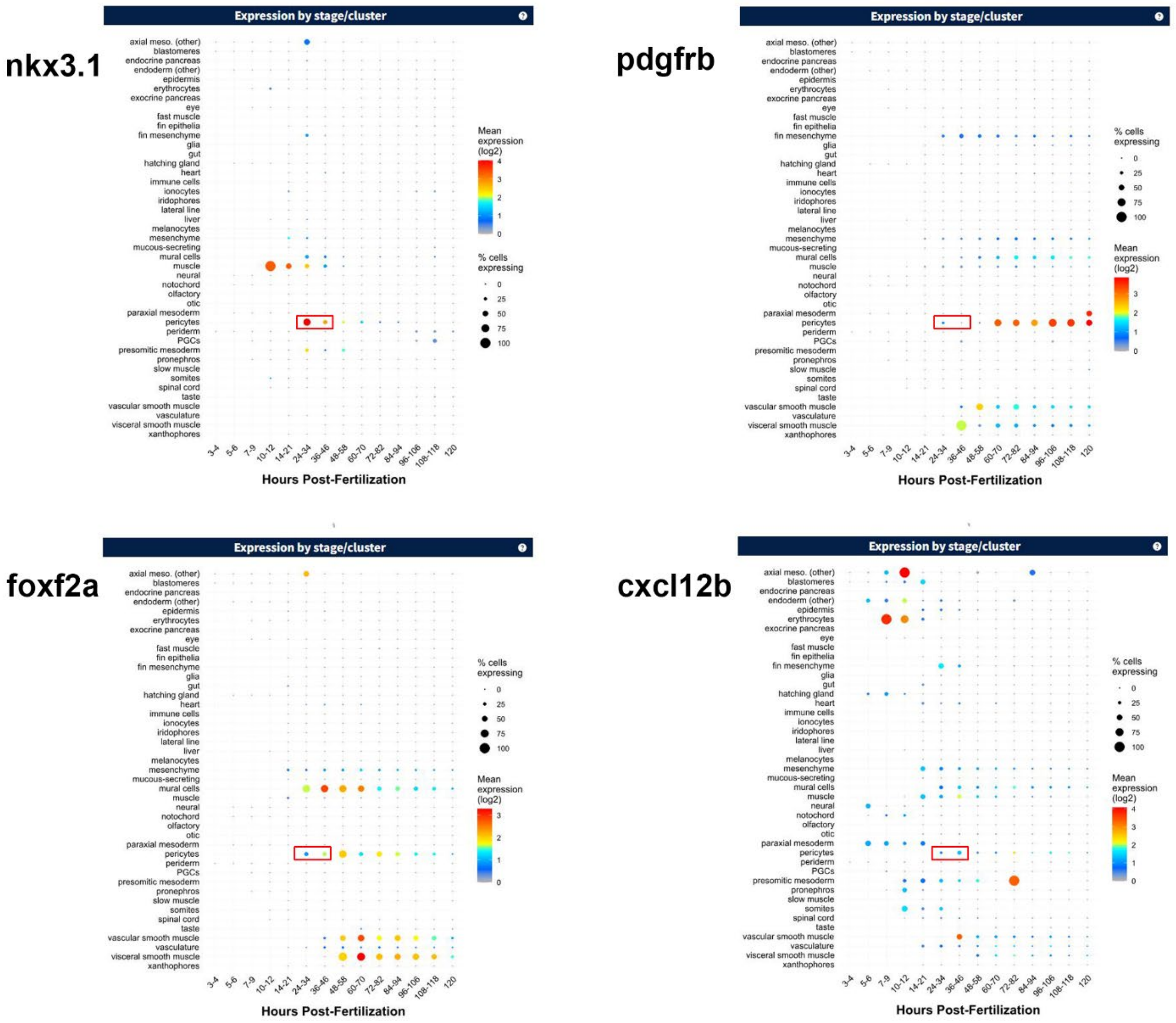


**Fig. S15: Expression of pericyte precursor genes in the Daniocell database** Output of searches for early pericyte markers showing the pericyte cluster expression at 24 hpf – 48 hpf of the indicated genes (red box).
